# Molluscan dorsal-ventral patterning relying on *bmp2/4* and *chordin* provides insights into spiralian development and bilaterian evolution

**DOI:** 10.1101/2020.08.11.245670

**Authors:** Sujian Tan, Pin Huan, Baozhong Liu

**Author notes:** These authors contributed equally to this work (Sujian Tan, Pin Huan). Corresponding author: Baozhong Liu, Key Laboratory of Experimental Marine Biology, Institute of Oceanology, Chinese Academy of Sciences, 7 Nanhai Road, Qingdao, 266071, China.

## Abstract

Although a conserved mechanism relying on *bmp2/4* and *chordin* is suggested in animal dorsal-ventral (DV) patterning, this mechanism has not been reported in spiralians, one of the three major clades of bilaterians. Studies on limited spiralian representatives have suggested markedly diverse DV patterning mechanisms, a considerable amount of which no longer deploy BMP signaling. Here, we showed that *bmp2/4* and *chordin* regulated DV patterning in the mollusk *Lottia goshimai*, which was predicted in spiralians but not reported before. In the context of the diverse reports in spiralians, it conversely represents a relatively unusual case. We then showed that *bmp2/4* and *chordin* coordinated to mediate signaling from the D-quadrant organizer to induce the DV axis, among which *chordin* transferred breakdown-of-symmetry information. Further investigations on the *L. goshimai* embryos with influenced DV patterning suggested roles of BMP signaling in regulating the localization of the blastopore and the organization of the nervous system, indicating a cooption of DV patterning and the transition of these key characteristics at the origin of bilaterians. These findings provide insights into the evolution of animal DV patterning, the unique development mode of spiralians driven by the D-quadrant organizer, and the evolution of bilaterian body plans.

---

The existence of a dorsal-ventral (DV) axis is a key characteristic in Bilateria. Generally, a conserved molecular logic, namely, the BMP ligand *bmp2/4* and its antagonist *chordin*, patterns the DV axis of bilaterians (1-6). It has been suggested that these two genes even pattern a body axis in nonbilaterian animal lineages (7, 8), indicating broad conservation. However, despite such conservation, the DV patterning mechanism exhibits a considerable degree of variation (9). In some cases, DV patterning no longer depends on *bmp2/4* and *chordin* (e.g., nematodes and ascidians (10, 11)). In two of the three major bilaterian clades, Ecdysozoa and Deuterostomia, such exceptional cases are considered lineage-specific characters since extensive evidence reveals *bmp2/4*-*chordin*-dependent mechanisms in their relatives (e.g., insects and vertebrates (9, 12)).

The situation in the other bilaterian clade, Spiralia, is very different. Unlike those in ecdysozoans and deuterostomes, the molecular mechanisms of spiralian DV patterning remain largely elusive. Moreover, current studies on several representative species have revealed quite diverse DV patterning mechanisms. These spiralians could use other BMP molecules (the leech annelid *Helobdella robusta*) (13) or even do not employ BMP signaling in DV patterning (the annelids *Capitella teleta, Chaetopterus pergamentaceus* and the mollusk *Crepidula fornicata*) (14-18). The roles of *bmp2/4* in DV patterning were proven in two spiralians (the platyhelminth *Dugesia japonica* and the mollusk *Tritia obsolete*) (19, 20), but it is unknown whether *bmp2/4* coordinates with *chordin* to induce correct DV patterning (as seen in most nonspiralian animals (9, 12)). This issue is important since *chordin* is suggested to be crucial in DV patterning (9), and this gene might have been lost from particular spiralian lineages (e.g., platyhelminths) (21). Together, although the ancestral *bmp2/4*-*chordin*-dependent DV patterning mechanism has been generally accepted for bilaterians (22), studies on six species spanning three spiralian phyla did not reveal such a mechanism (Fig. 1a).

**Fig. 1.**
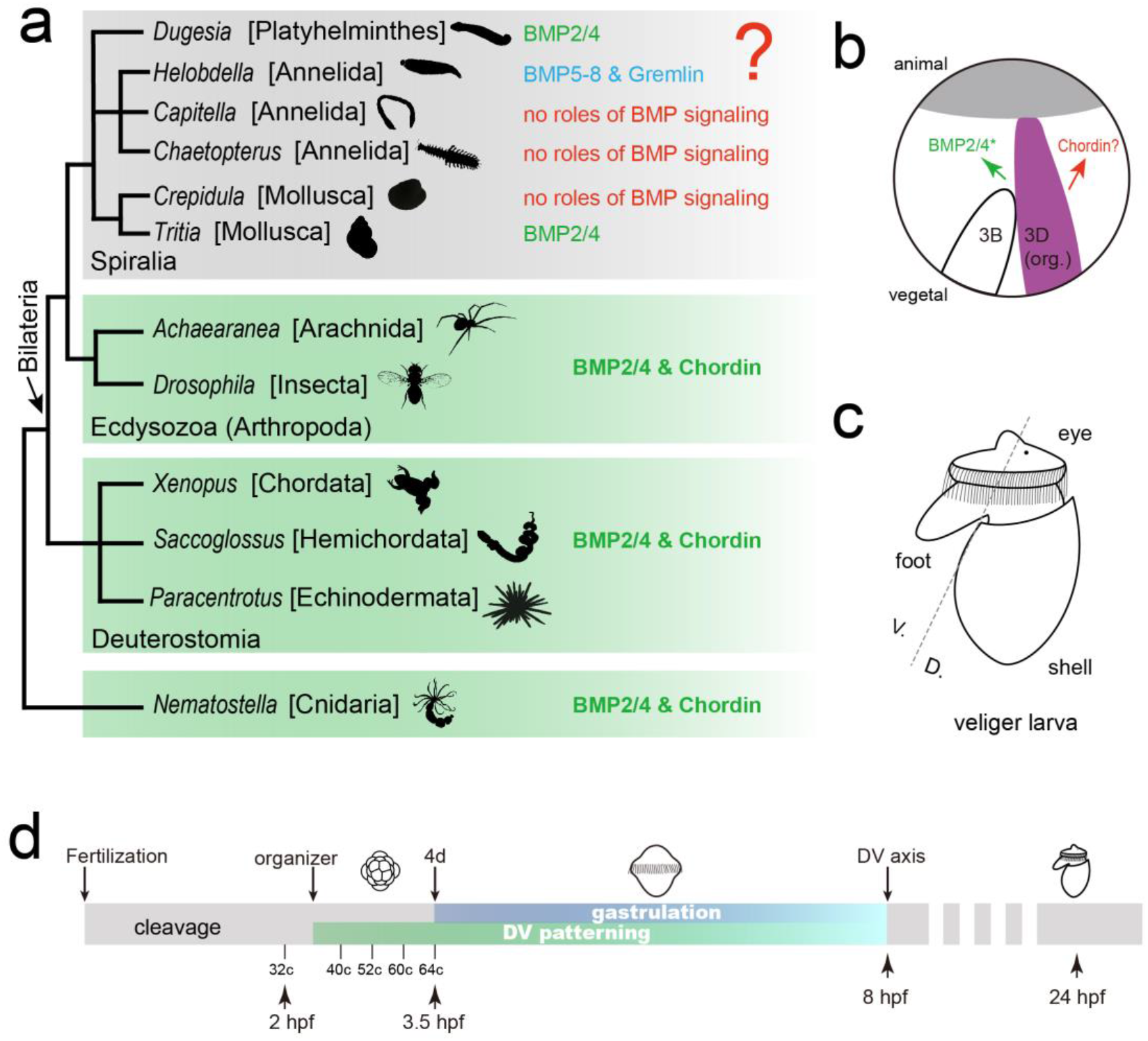
Mollusks represent ideal systems to understand the evolution of animal DV patterning and spiralian organizer function. a. Conserved roles of *bmp2/4* and *chordin* in animal body patterning. In most major animal clades, *bmp2/4* and *chordin* determine a body axis (it is the DV axis for bilaterians). For spiralians, however, the knowledge is elusive, and some results argue against this conserved mechanism (blue and red letters). The diagrams of representative animals are derived from PhyloPic (http://phylopic.org/) and Wikipedia (https://www.wikipedia.org) licensed under CC BY 3.0. **b-c. Unique spiralian developmental mode relying on a D-quadrant organizer**. Panel **b** shows that after its formation, the organizer (3D) activates BMP signaling by regulating *bmp2/4* (green letters); however, whether *chordin* is involved in this process remains unknown (red letters). The asterisk indicates that this mechanism requires certifications in more species. Panel **c** shows a veliger larva of gastropod mollusks, and the processes regulated by the organizer are highlighted: DV patterning (generally indicated by the dashed line) and the formation of marked larval organs. **d. Early development of the gastropod mollusk *L. goshimai* (at 25°C) emphasizing DV patterning events**. Organizer formation at the middle 32-cell (32c) stage marks the beginning of DV patterning, which is largely coupled with gastrulation since the formation of the 4d blastomere at the 64-cell stage. A DV axis was well established at 8 hpf, and a characteristic veliger larva is developed at approximately 24 hpf.

Despite the suggested diversity at the molecular level, spiralian DV patterning actually shows considerable conservation at the cellular level. In spiralian lineages such as annelids, mollusks and nemerteans, the DV axis is induced by a D-quadrant organizer, referring to a special blastomere that regulates the development of the whole embryo (e.g., 3D or 4d, according to the nomenclature for describing spiral cleavage; see Fig. 1b and c) (23-25). In fact, the involvement of the D-quadrant organizer, in parallel with several other characteristics conserved in this clade but infrequently observed in other clades (e.g., spiral cleavage), is suggested to be the most important characteristic comprising the unique developmental mode of spiralians (23, 26-29). Although the developmental functions of spiralian D-quadrant organizer have been well determined, only limited knowledge is known regarding how it functions at the molecular level (30-33). In this context, investigating the molecular mechanisms of DV patterning would be an essential aspect to decipher the organizer function for spiralians, given the conserved role of organizer in inducing the DV axis.

Interestingly, a link was recently established between organizer and the DV patterning gene *bmp2/4*. A pioneering report proved that *bmp2/4* mediated organizer signaling and regulated DV patterning in the gastropod mollusk *Tritia* (20), explaining the conserved organizer function to induce DV patterning. Despite this essential process, open questions still exist. First, it is unknown whether such *bmp2/4*-dependent organizer function also exists in other spiralian lineages. Investigations on the prevalence of such a mechanism are necessary given the suggested diversity in spiralian DV patterning mechanisms at the molecular level, as mentioned above. Moreover, the dynamics of organizer signaling could differ significantly between unequal cleavers and equal cleavers (even from the same class, e.g., the snail *Tritia* and the limpet *Tectura*) (30, 31), adding to the question of whether they would utilize the same molecule to execute organizer function. Second, and more importantly, it should be explored whether a BMP antagonist is involved in organizer function. This question should be clarified since the crucial node in DV patterning (i.e., organizer function) is not the BMP ligand itself but the gradient of BMP signaling along the presumed DV axis (12). Such a spatial distribution of BMP signaling is largely determined by extracellular BMP regulators such as *chordin* (12, 34). In fact, the most important gene in DV patterning is proposed to be *chordin* but not *bmp2/4*. Restricted *chordin* expression is suggested to be sufficient to determine a BMP signaling gradient irrespective of the expression patterns of *bmp2/4* (9). Thus, despite knowing the involvement of *bmp2/4* in organizer function (in *Tritia*, still requiring investigations in additional species), a key question that follows is whether and how the organizer induces the BMP signaling gradient. Given the conserved roles of *chordin* as a major BMP antagonist gene in animal DV patterning, this gene could be the primary candidate to address this issue.

Mollusks emerge as ideal systems to clarify the abovementioned questions, i.e., whether *bmp2/4* and *chordin* function in DV patterning of spiralians and the relationship between the two genes and the D-quadrant organizer. The organizer function to induce the DV axis has been well investigated in mollusks (35-37). Although the roles of BMP signaling in DV patterning have not been revealed in *Crepidula* (14), they have been demonstrated in *Tritia* (20). Our results using a small molecule BMP inhibitor also support this notion (in the bivalve *Crassostrea gigas*) (38). Moreover, unlike some spiralians that likely lack the *chordin* gene, the gene was identified in mollusks (21). We further showed that *bmp2/4* and *chordin* were expressed on opposite sides along the DV axis of the *Crassostrea* embryo, indicating roles in DV patterning (39). Here, we investigated the roles of *bmp2/4* and *chordin* in the gastropod mollusk *Lottia goshimai*. We confirmed that the two genes both regulated DV patterning and participated in organizer function. By examining embryos with influenced DV patterning, we further revealed evidence suggesting the profound developmental effects of stereotype cleavage and the regulatory roles of BMP signaling in the localization of blastopore and the organization of the nervous system. These findings provide insights into the unique developmental mode of spiralians and the evolution of bilaterian body plans.

## Results

### bmp2/4 and chordin mediate organizer signaling and determine the BMP signaling gradient

Both *bmp2/4* and *chordin* were retrieved from the developmental transcriptome of *L. goshimai*. Given that molluscan DV patterning relies on the D-quadrant organizer, we first investigated the expression of *bmp2/4* and *chordin* around the time of organizer formation (from the 16-cell to 64-cell stage; see supplemental text for details about the *L. goshimai* organizer (3D)). At the same time, the dynamics of BMP signaling were explored by tracing the key signal transducer phosphorylated Smad1/5/8 (pSmad1/5/8). During the period investigated, *bmp2/4* was expressed in a generally radial pattern with minor changes (supplemental Fig. S2a-d). In contrast, we found a strong correlation among *chordin* mRNA expression, BMP signaling and the organizer (Fig. 2a-h and supplemental Fig. S2). In brief, sequential developmental events were observed in this period: 1) organizer formation (32-cell stage), 2) activation of universal BMP signaling (late 32-cell stage, Fig. 2f, 3) transition of *chordin* expression into an asymmetrical pattern (32-to-40-cell stage, Fig. 2c, and 4) transition of BMP signaling into an asymmetrical pattern (52-/60-cell stage, Fig. 2h). In the 60-cell embryo, the cells adjacent to the organizer showed strong BMP signaling, while only weak signaling was detected in the cells distal to the organizer (Fig. 2h and supplemental Fig. S2). This distribution pattern is comparable to the BMP signaling gradient along the DV axis in many animals (e.g., *Drosophila* and sea urchin) (40, 41); we thus refer to it as the BMP signaling gradient, although it does not exhibit an exact gradient pattern likely related to the small cell numbers and large cell volumes in *L. goshimai* embryos. Since the direction of this gradient was across the 3D and 3B blastomeres (Fig. 2h) that generally coincided with the presumptive DV axis (23, 27), this BMP signaling gradient marked a molecular DV axis prior to the morphologically detectable DV axis.

**Fig. 2.**
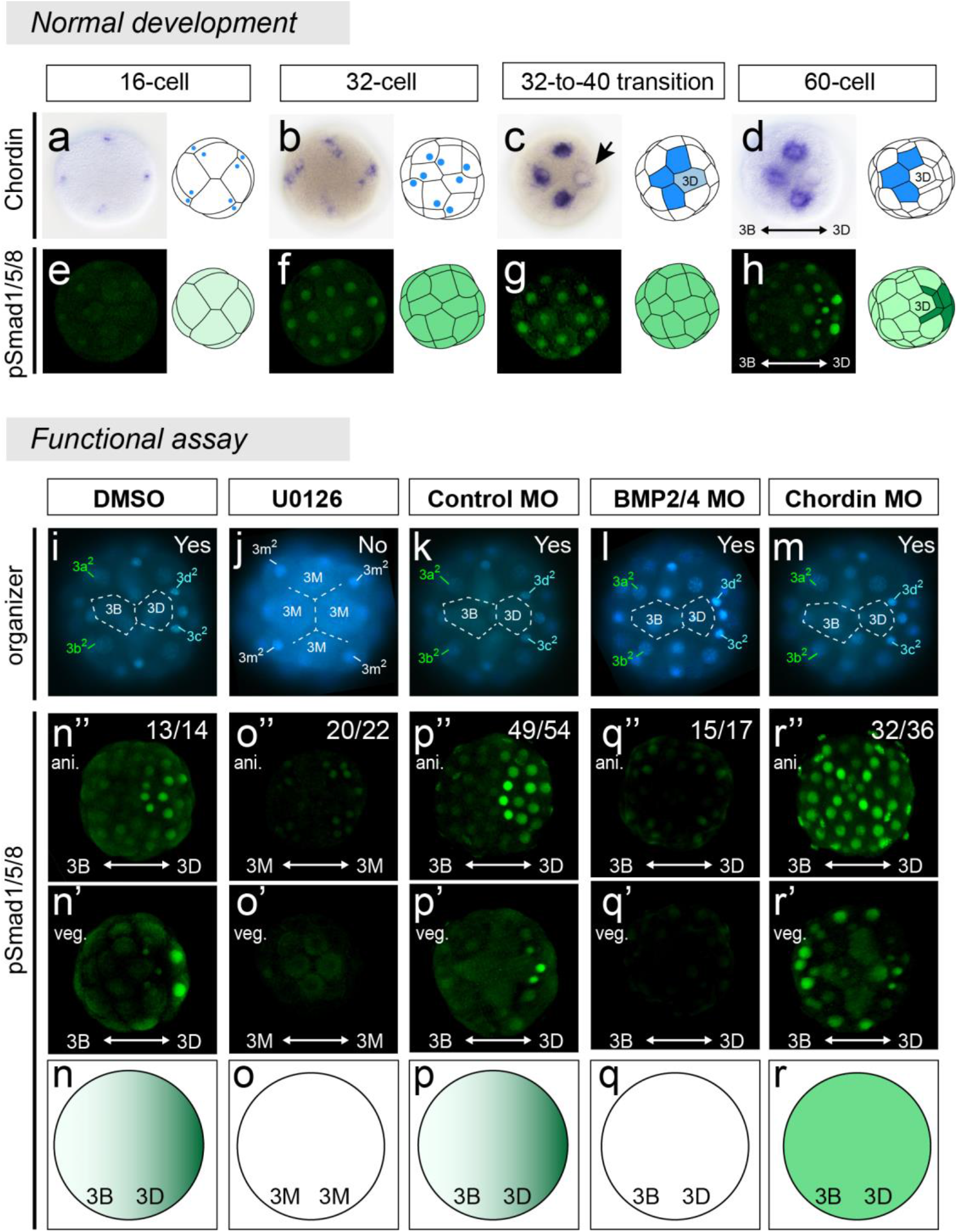
Regulatory relationships among the organizer, *bmp2/4* and *chordin*. a-h. Vegetal views, *chordin* expression (**a**-**d**) and the state of BMP signaling that is indicated by phosphorylated Smad1/5/8 (pSmad1/5/8) staining (**e-h**, confocal projections) from the 16- to 60-cell stage. The arrow in **c** indicates weakened *chordin* expression in the organizer (3D) at the late 32-cell stage. More details are provided in supplemental Fig. S2. **i-r**. the states of organizer **(i-m)** and BMP signaling **(n-r)** under different manipulations, all samples at the 60- or 63-cell stage. In **i-m**, the organizer is identifiable based on the characteristic 4-cell arrangement at the vegetal pole (see supplemental Fig. S1a), and whether an organizer was formed is indicated by “yes” or “no” in the panels. **n-r**. Diagrams showing pSmad1/5/8 staining along the 3B-3D axis (lateral views with the animal pole to the top). Both animal (ani., **n’-r’**) and vegetal (veg., **n’’-r’’**) views are shown.

The correlations among organizer, *chordin* expression and BMP signaling suggest regulatory relationships. We first confirmed that organizer formation triggered BMP signaling. When organizer formation was inhibited by the MAPK inhibitor U0126 (as described previously (30)) (Fig. 2j), the activation of BMP signaling was prevented (Fig. 2o). We then found that such activation of BMP signaling was mostly mediated by *bmp2/4* because injecting an antisense *bmp2/4* morpholino (MO) largely eliminated pSmad1/5/8 staining (Fig. 2q), while it did not influence organizer formation (Fig. 2l). The regulation of *bmp2/4* function by the organizer should be at the posttranscriptional level, since no significant change in *bmp2/4* mRNA expression was detected before and after organizer formation (supplemental Fig. S2a-d) or after U0126 treatment (supplemental Fig. S3h). However, the BMP signaling activated by the organizer only showed a universal distribution. We revealed that *chordin* was required to transit this universal distribution in a gradient manner. When *chordin* was inhibited by injecting an antisense MO, no gradient formed, and universal BMP signaling was sustained in subsequent developmental stages (till at least 64-cell stage, Fig. 2r), despite the normally formed organizer (Fig. 2m). We found that *chordin* expression was also regulated by the organizer. When organizer formation was inhibited, symmetrical *chordin* expression was no longer interrupted (supplemental Fig. S3).

Based on these results, we conclude the regulatory relationships among organizer, *bmp2/4, chordin*, and BMP signaling in *L. goshimai* (Fig. 3). In brief, after its formation, the organizer triggers *bmp2/4* (green arrows in Fig. 3b), which induces universal BMP signaling activities (Fig. 3c). Shortly afterwards, the organizer regulates *chordin* expression to become an asymmetrical pattern, which modulates BMP signaling to form a gradient along the presumptive DV axis (Fig. 3d). Taken together, under the regulation of the organizer, *bmp2/4* and *chordin* coordinate to generate the correct BMP signaling gradient: *bmp2/4* activates signaling, and *chordin* determines the spatial distribution (gradient) of signaling. From this point of view, *chordin* is the key molecule to transfer the breakdown-of-symmetry signal from the organizer to form the secondary body axis.

**Fig. 3.**
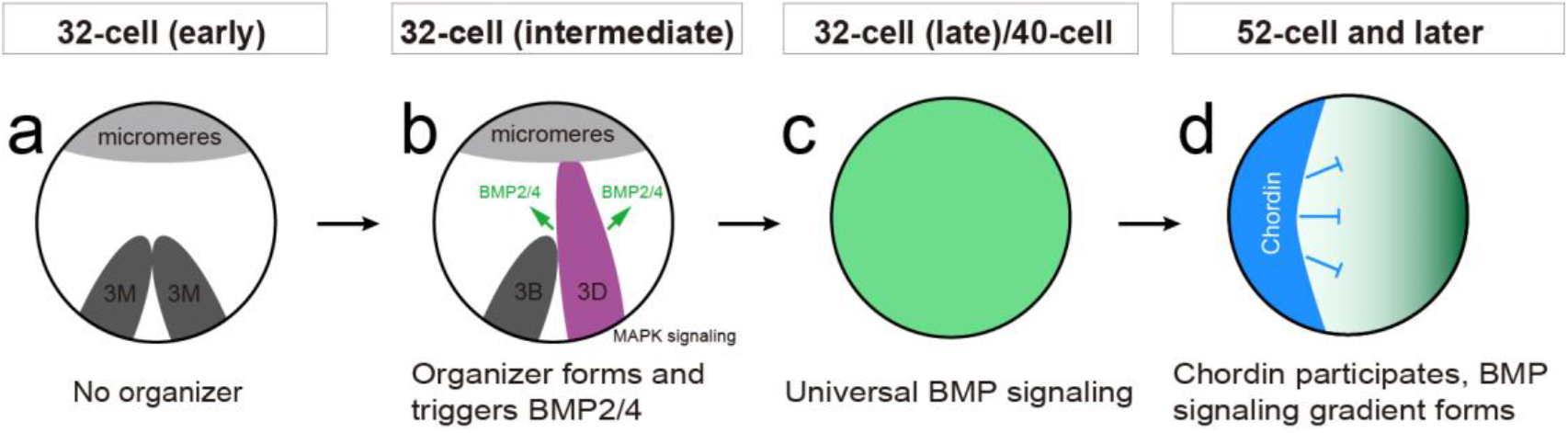
A hypothesis assuming regulatory relationships among organizer, *bmp2/4* and *chordin*. *bmp2/4* and *chordin* coordinate to generate the correct BMP signaling gradient to regulate DV patterning, which was regulated by the organizer. When the 32-cell embryo initially forms, the four macromeres (3M) are equivalent (**a**). One of them is subsequently induced to be the organizer (3D) due to the establishment of direct contacts with micromeres at the animal pole (42); MAPK signaling is then activated in this blastomere (30) (**b**). The organizer triggers *bmp2/4* (green arrows in **b**), which induces universal BMP signaling activities (**c**), and then it also regulates *chordin* expression to further transform BMP signaling into a gradient pattern (**d**). See other information in the text.

### Normal DV patterning and expression of bmp2/4 and chordin

Since the formation of the BMP signaling gradient, DV patterning of *L. goshimai* began, which was largely coupled with the characteristic epibolic gastrulation in this gastropod lineage (e.g., *Patella* (43, 44)) (Fig. 1d). A DV axis was well established at 8 hpf, reflected by the development of a shell field on the dorsal side and that of the ventral plate and blastopore on the ventral side (Fig. 4a-j). Since these structures were morphologically detectable at relatively late developmental stages, we investigated the expression of several marker genes to explore the details of *L. goshimai* DV patterning. The expression of blastopore marker genes (*brachyury* and *foxa*) was asymmetrical along the DV (3B-3D) axis since the very beginning of DV patterning (the 60-cell stage, ∼3.2 hpf) (supplemental Fig. S4a-c). However, given that the blastopore only represented a small portion of embryonic cells (Fig. 4h-j), we focused on the shell field and the ventral plate that occupied most of the area of the dorsal/ventral surface in the 8-hpf embryo (Fig. 4b-g).

**Fig. 4.**
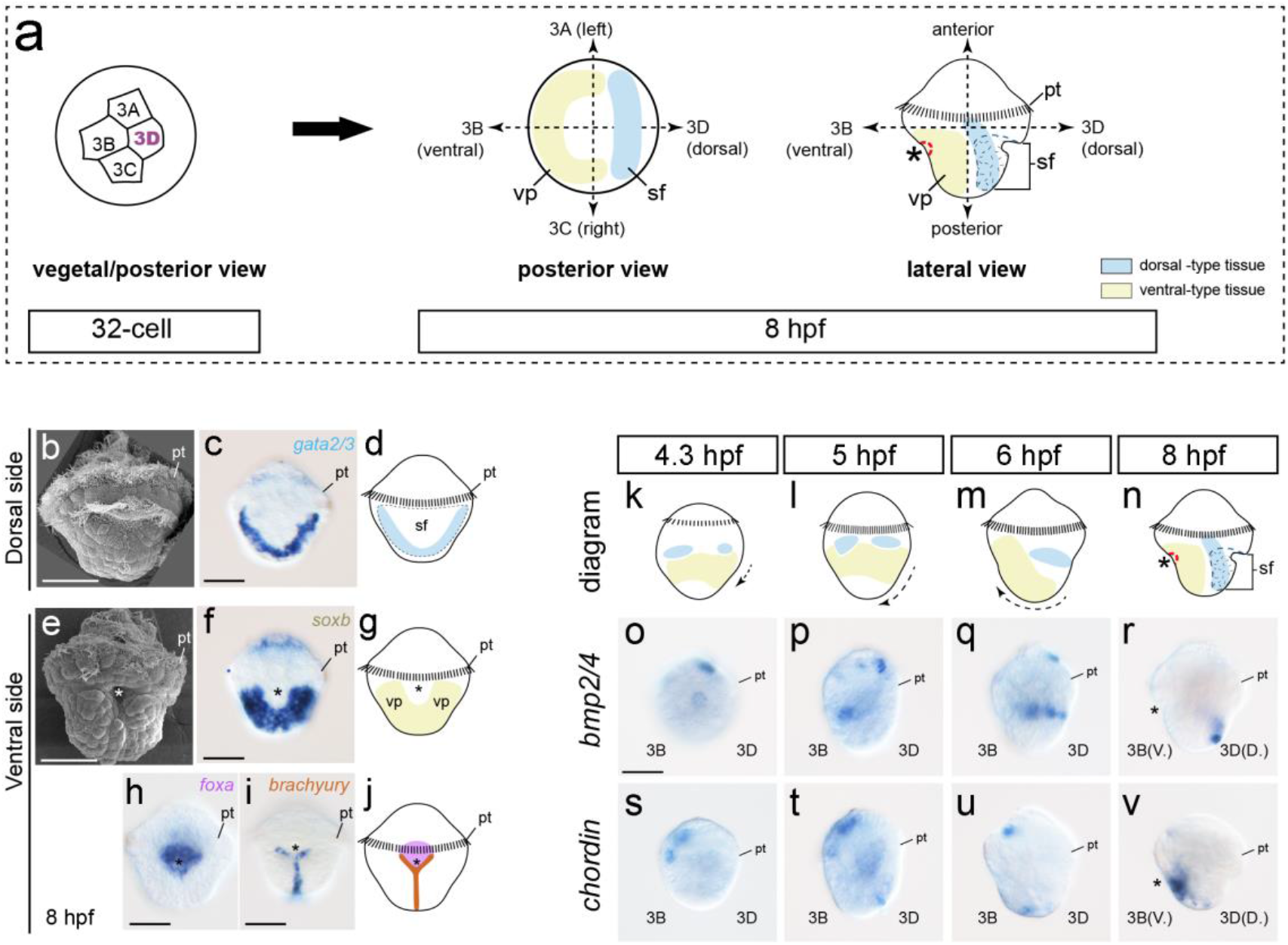
Normal DV patterning of *L. goshimai* and *bmp2/4* and *chordin* expression during this period. The diagrams in **a** show that the DV axis at 8 hpf corresponds to the 3B-3D axis at the 32-cell stage. **b-j**. Morphological characters of the 8-hpf embryos. On the dorsal side, a shell field (sf) is discriminable, the out edge of which expresses the marker gene *gata2/3* (**b-d**). On the ventral side (**e**), two structures are developed, including the ventral plate (vp) expressing *soxb* (**f** and **g**) and the blastopore (asterisks) marked by *foxa* expression as well as a part of *brachyury* expression (**h-j**). **k-v**. Normal expression of *bmp2/4* and *chordin* during DV patterning, lateral views anterior to the top. Diagrams in **k-n** show the movements of dorsal- and ventral-type tissues during DV patterning based on the dynamics of *gata2/3* and *soxb* expression (supplemental Fig. S4). The asterisks indicate the blastopore. pt, prototroch. Bars represent 50 μm.

Two marker genes were investigated, including *gata2/3*, which was expressed in the outer edge of the shell field (as in another mollusk (45), Fig. 4c), and *soxb*, which was universally expressed in the ventral plate (46) (Fig. 4f). Expression of both genes indicated that at the initial phase of DV patterning (4.3-5 hpf), the anlages of the shell field and ventral plate were aligned generally along the anterior-posterior (AP) axis (Fig. 4k, l and supplemental Fig. S4d-i), themselves organized in almost circular patterns (supplemental Fig. S4d’-i’). In subsequent development, the two tissues moved with the epibolic gastrulation and were ultimately located dorsally and ventrally at 8 hpf (Fig. 4k-n and supplemental Fig. S4j-o). During this period, posttrochal expression of *bmp2/4* and *chordin* correlated with the dynamics of the two tissues (Fig. 4o-v), and they were also distributed on the dorsal or ventral side at 8 hpf (Fig. 4r and v). This correlation indicates the roles of the two genes in DV patterning.

### Radialized early development: DV patterning relying on bmp2/4 and chordin

When inhibiting either *bmp2/4* or *chordin* by injection with specific MOs, DV patterning of *L. goshimai* was largely inhibited. As shown in Fig. 5, at 6 hpf, compared to the asymmetrical gene expression in normal embryos (*gata2/3, soxb, bmp2/4* and *chordin*), the embryos with *bmp2/4* or *chordin* knockdown showed generally radial expression (Fig. 5f-i, k-n) (despite significant differences between the two phenotypes, see more details in supplemental Fig. S5). Treatment of early embryos with 0.5 µg/ml recombinant human BMP4 protein (rhBMP4) also generated a similar radialized phenotype (supplemental Fig. S6h-m). The absence of a DV axis in these radialized phenotypes suggests that the DV patterning of *L. goshimai* was inhibited when *bmp2/4* or *chordin* was knocked down, revealing the essential roles of the two genes.

**Fig. 5.**
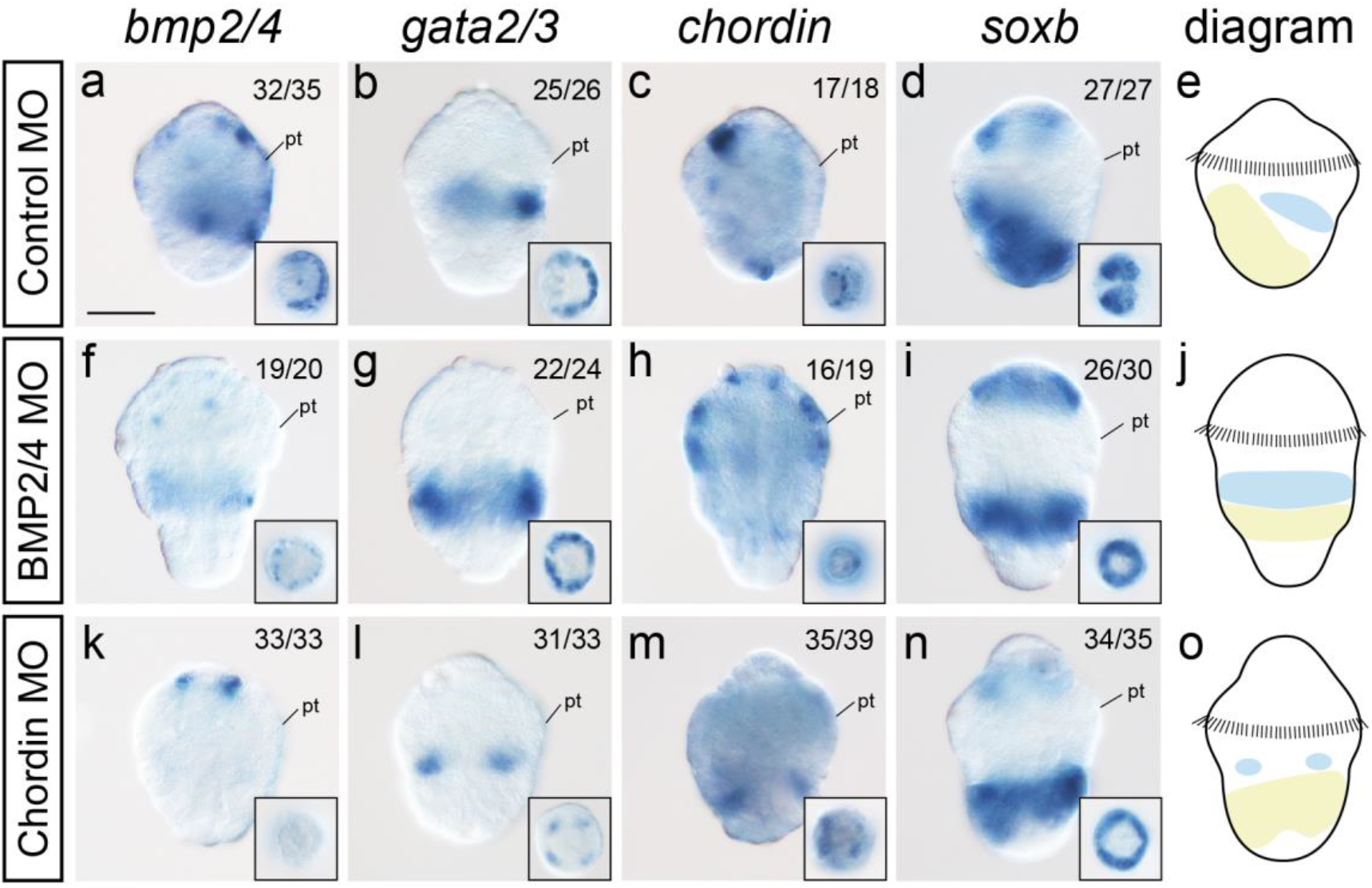
*bmp2/4* and *chordin* knockdown phenotypes at 6 hpf. All panels are posterior views anterior to the top, and the inserts show posterior views. The diagrams are derived from the expression patterns of *gata2/3* (dorsal-type tissues) and *soxb* (ventral-type tissues). After knockdown of *bmp2/4* or *chordin*, these genes generally show radial expression. The dorsal- and ventral-type tissues were aligned along the AP axis in the influenced embryos. Notably, the expression of *chordin* and *soxb* in the *chordin*-knockdown embryo actually showed minor asymmetry at this stage (**m** and **n**). However, despite this, we think these embryos were comparable with the radialized phenotypes caused by *bmp2/4* knockdown (**f-i**) or rhBMP4 protein treatment (supplemental Fig. S6h-m). See more details in supplemental Fig. S5. pt, prototroch. Bars represent 50 μm.

The expression levels of the marker genes in the *bmp2/4*- or *chordin*-knockdown embryos did not show a certain trend of changes, indicating inhibitory or promoting effects of BMP signaling. Although it seemed that BMP signaling inhibited *gata2/3* expression (Fig. 5g and l), it was difficult to conclude whether the expression levels of *soxb* were up- or downregulated in any groups (based on ISH; Fig. 5i and n). Similarly, no obvious trends of expression change were observed for *bmp2/4* or *chordin* themselves (Fig. 5f, h, k and m). Despite such uncertainty, a common characteristic of the knockdown embryos was that the dorsal-type (*gata2/3*-positive) and ventral-type tissues (*soxb*-positive) were distributed along the AP axis (Fig. 5j and o), themselves showing radial organizations (inserts in Fig. 5f-i, k-n), reminiscent of the embryonic body pattern at the initial phase of DV patterning (Fig. 4k and l). Therefore, these knockdown phenotypes seem to indicate that the major role of the DV patterning signal (manifested by the BMP signaling gradient of early embryos) is to drive the dorsal/ventral-type tissues, which are anteriorly-posteriorly distributed initially, to move to their destined locations. When the BMP signaling gradient was eliminated, either by inhibition of signaling (*bmp2/4* knockdown) or generation of a universal distribution (*chordin* knockdown), the movement of the tissues was likely interrupted, producing the arrest of the “initial state” of the embryonic body plan.

Last, although a major role of BMP signaling is suggested to regulate cell movements, its roles in regulating cell specification are also suggested. Despite the largely circular organization of early *gata2/3* and *soxb* expression, asymmetry was detectable at early developmental stages (supplemental Fig. S4d’-i’). This result indicates that the initial dorsal/ventral-type tissues were not fully radial as in the influenced embryos and that BMP signaling indeed contributed to the specification of these tissues. Taken together, our results indicate that the DV patterning signal (the BMP signaling gradient) plays two roles in DV patterning: 1) it causes polarized specification of embryonic cells, though related tissues are still organized in a largely circular pattern, and 2) more importantly, it drives these tissues, which are distributed along the AP axis initially, to move to their destined locations along the DV axis.

### Reemerged asymmetrical development and posteriorized blastopore in late embryos

A close look into the *chordin*-knockdown embryos actually revealed minor asymmetry (Fig. 5m and n), and we found that this asymmetry was significantly amplified in subsequent development. Somewhat unexpectedly, minor asymmetry also remerged in the *bmp2/4*-knockdown embryos. For clarity, in the following text, we will describe the orientation of the manipulated embryos based on the locations of 3Q blastomeres (e.g., the 3B or 3D side) since normal DV patterning was influenced. In normal embryos, the 3B and 3D sides correspond to the ventral and dorsal sides, respectively (Fig. 4a).

At 8 hpf, the normal embryos had a well-developed DV axis (Fig. 6a-f). Due to the influence of DV patterning, the *bmp2/4*-knockdown embryos largely retained radial development (Fig. 6g-l and supplemental Fig. S7f-j). Only minor asymmetry was detected (e.g., polarized *chordin* expression; see Fig. 6i and supplemental Fig. S7i); we could not determine the direction of the asymmetry. In contrast, asymmetrical development in the *chordin*-knockdown embryos was much more evident (Fig. 6m-r and supplemental Fig. S7k-o). Such asymmetry occurred along the 3B-3D axis (see supplemental Fig. S8 for details on the orientation of the manipulated embryos). In particular, in the posterior part of the embryo, the 3D side exhibited much greater development than the 3B side (red arrows in Fig. 6m-q), which showed *soxb* expression marking ventral-type tissues (Fig. 6o). The posterior tissues on the 3D side were further divided into two bilateral lobes to make the embryo exhibit a pseudotwin phenotype (supplemental Fig. S7n and o). In a rare case, such a pseudotwin phenotype even developed duplicated larval shells (supplemental Fig. S9).

**Fig. 6.**
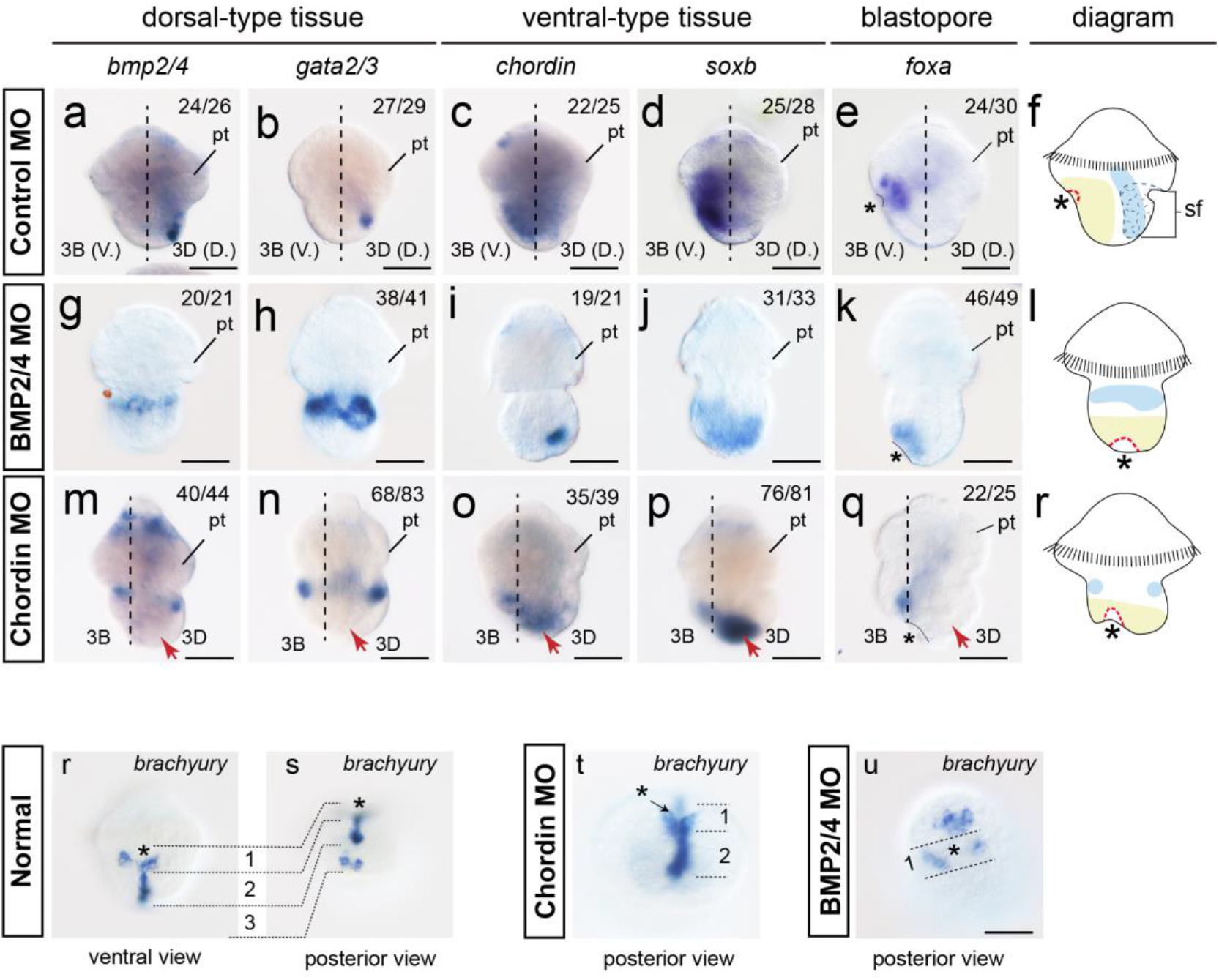
*bmp2/4* and *chordin* knockdown phenotypes at 8 hpf. Panels **a-r** show lateral views with the 3B side to the left (when discriminable, indicated by dashed lines). The diagrams in **f, l** and **r** show the body plans derived from the marker gene expression patterns shown in other panels. Radial development was largely sustained in the *bmp2/4*-knockdown embryos (**g-k**), despite polarized expression of *chordin* (**i**). However, evident asymmetry was observed for *chordin*-knockdown embryos (red arrows in **m**-**q**), although the dorsal- and ventral-type tissues were still generally distributed along the AP axis as the *bmp2/4*-knockdown embryos (**r** and **l**, compare them to the 6-hpf embryos in Fig. 5). Expression of the blastopore marker *foxa* indicates that locations of the blastopore (asterisks) are very different in normal embryos (**e**, ventral) and manipulated embryos (**k** and **q**, posterior). See more details in supplemental Fig. S7. Panels **r**-**u** shows *brachyury* expression at 8 hpf. At this stage, normal *brachyury* expression comprises three parts (indicated by numbers in **r** and **s**; see the text for more information). After gene knockdown, expression part 1 could be discriminated in both types of embryos (**t** and **u**), while expression part 2 was only observed in the *chordin*-knockdown embryo (**k**). The bars represent 50 μm.

The blastopore was also formed in 8-hpf embryos. Notably, we found that the blastopore was posteriorized in both types of embryos (supplemental Fig. S7f and k), showing sharp contrast with the normal blastopore formed ventrally (supplemental Fig. S7a). The posteriorized blastopore was confirmed by the expression of the blastopore marker gene *foxa* (Fig. 6k and q, compared to the normal ventral expression shown in Fig. 6e). Such posteriorized blastopore is consistent with the fact that the influenced larval body plan exhibited tissues aligned along the AP axis (Fig. 6l and r). In *L. goshimai*, the blastopore forms as a consequence of extensive cell movements during gastrulation (similar to *Patella* (43, 44)). Therefore, the changes in blastopore location in these manipulated embryos suggest altered cell movements during gastrulation, supporting our speculation that an essential role of the DV patterning signal is to regulate cell movement.

Since the posteriorized blastopore may have essential evolutionary implications, we sought to explore whether there was molecular evidence to indicate the developmental capacity of blastoporal cells, given that it was difficult to directly trace the development of the blastopore due to seriously disturbed development. To this end, we investigated the expression of another blastopore marker gene, *brachyury*, in the manipulated embryos. Similar to another gastropod (44), the normal *brachyury* expression of *L. goshimai* comprised three parts at 8 hpf (indicated by numbers in Fig. 6r and s): 1) the posterior edge of blastopore, which showed a V shape and would contribute to the formation of larval mouth; 2) the ventral midline; and 3) posterior expression without determined fates. We found that in both *bmp2/4*- and *chordin*-knockdown embryos, although *brachyury* expression changed considerably, a V-shaped expression pattern was still discriminable in blastoporal cells (number “1” in Fig. 6t and u). This result indicates that despite the posteriorized localizations, the blastoporal cells seemed to still retain the developmental potential of the larval mouth in the manipulated embryos.

### Effects of BMP signaling on neurogenesis in L. goshimai

There has been extensive evidence indicating the roles of the molluscan organizer in eye development, with some inconsistency among studies (31, 47-49). A recent study placed eye development into the context of neurogenesis and suggested a positive role of BMP signaling in neurogenesis of the gastropod mollusk *Tritia* (20). This effect, however, is opposite to nearly all other animals (50). We therefore investigated the relationship between BMP signaling and eye development/neurogenesis in *L. goshimai*. We found that, consistent with previous reports (20, 47, 48), BMP signaling (i.e., the inductive signals from the organizer) promoted eye development in *L. goshimai*: no eye formed after *bmp2/4* knockdown, and extra eyes formed under hyperactive BMP signaling when *chordin* was inhibited (Fig. 7a-c). rhBMP4 treatment also generated similar phenotypes (although a dose-dependent effect was indicated; see details in supplemental Fig. S6o’), which are comparable to that of a previous study (20).

**Fig. 7.**
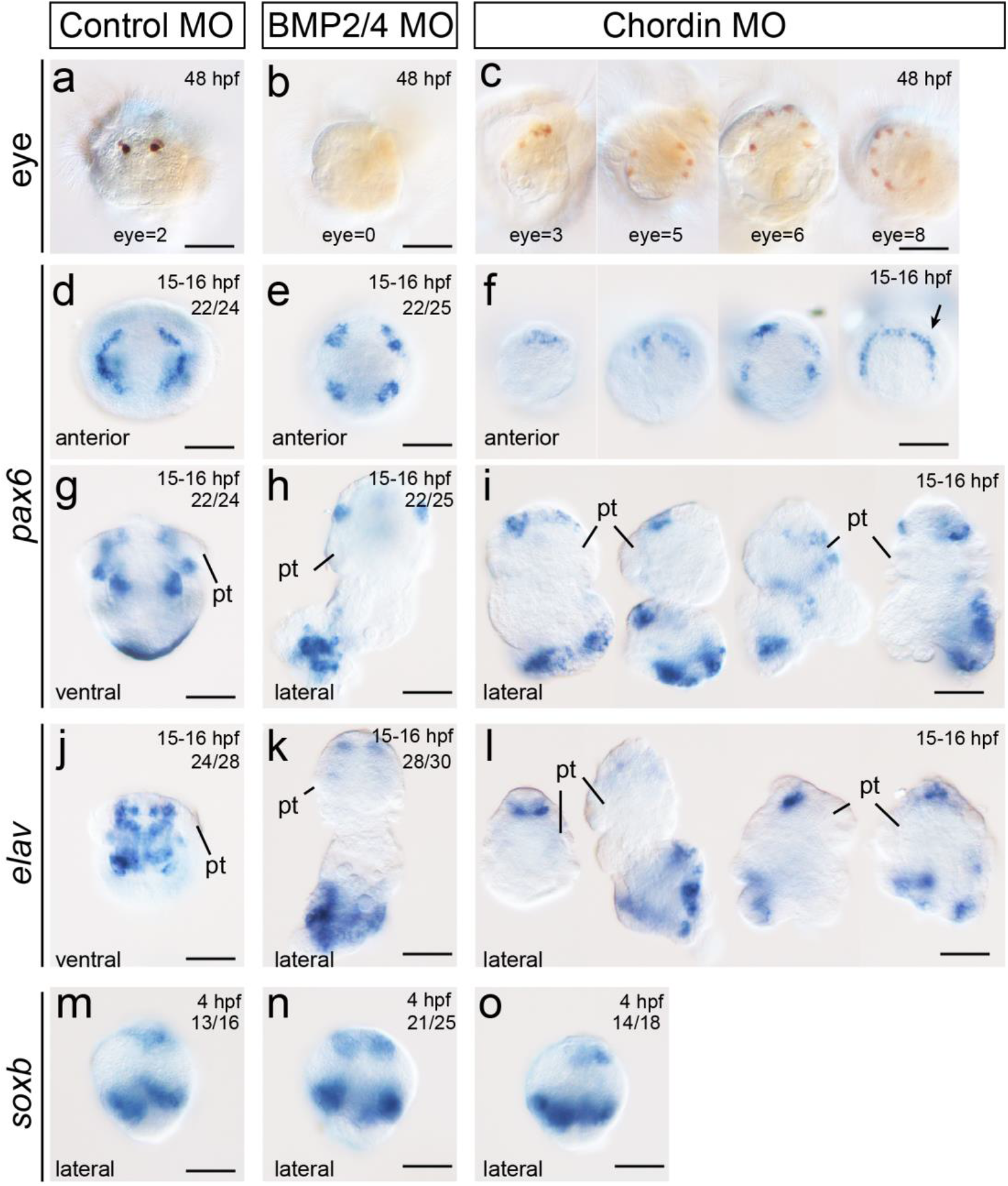
Effects of BMP signaling on eye formation and neurogenesis. **a-c.** Anterior views showing the eyes of 48-hpf larvae; a summary is shown in supplemental Fig. S10a. **d-o**. Expression of the neural markers *pax6* and *elav* in larvae (15-16 hpf) and *soxb* expression in early gastrulae (4 hpf). Since *pax6* and *elav* expression after *chordin* knockdown shows relatively high levels of heterogeneity, representative larval phenotypes are provided (**f, i** and **l**), and it is not possible to provide the number of individuals. Pretrochal *pax6* expression shows apparent correlations with eye distribution (compare **c** and **f**); however, there is no one-to-one relationship between the two panels. pt, prototroch. The bars represent 50 μm. More details on early *soxb* expression are provided in supplemental Fig. S10b-g.

However, subsequent analyses of neural marker genes did not support the proposal (20) that BMP signaling promotes neurogenesis. Although *chordin* inhibition expanded the expression of the neural patterning gene *pax6* in a portion of larvae (the arrow in Fig. 7f), *pax6* expression in the *chordin*-knockdown larvae exhibited considerable heterogeneity (Fig. 7f, i); thus, it was difficult to conclude a general pattern. Moreover, after *bmp2/4* knockdown, *pax6* expression did not show the expected downregulation (Fig. 7e, h). Therefore, *pax6* expression in the manipulated larvae did not suggest whether neurogenesis was promoted or inhibited by BMP signaling. Although expanded *pax6* expression in BMP4-treated *Tritia* larvae is considered an indicator of promoted neurogenesis (20), this result can also be interpreted to reflect the development of extra eyes given that *Tritia pax6* expression was mostly detected in the pretrochal region at the stage examined (20) and that we found that *pax6* expression in the pretrochal region showed an apparent correlation with the distributions of larval eyes (when BMP signaling was activated, compare Fig. 7c and f).

Given that *pax6* might not be an appropriate marker for overall neurogenesis (it might only contribute to the development of subpopulations of neural tissues) and that the conserved roles of BMP signaling in neurogenesis may be detectable only in the early phase of neurogenesis (20), we analyzed two additional marker genes. Among them, *elav* is a universal neuron marker (51), and its expression pattern was proven to coincide with the distribution of neural tissues in a mollusk (52). The other marker gene, *soxb*, plays essential roles in the early phase of neurogenesis (53). *elav* expression was similar to that of *pax6* (Fig. 7j-l), and it is difficult to conclude whether neurogenesis is inhibited or promoted in any group. For *soxb*, we focused on its expression at the stage when gastrulation was just beginning (4 hpf). Neurogenesis should be in the early phase at this stage (featuring processes, such as definition of the neuroectoderm and commitment of neural stem cells). We found that although *soxb* expression indeed changed after inhibition of *bmp2/4* or *chordin* (supplemental Fig. S10b-g), it was not highest after *bmp2/4* knockdown or lowest after *chordin* knockdown (Fig. 7m-o), similar to the expression at 6 hpf. These results also did not indicate whether BMP signaling inhibits or promotes neurogenesis in *L. goshimai*.

Instead of indicating positive or inhibitory effects, our results suggest that BMP signaling seems to be irrelevant to neurogenesis per se but affects the organization of the nervous system. As revealed by both *pax6* and *elav* expression, a common phenotype after *bmp2/4* or *chordin* knockdown was the loss of featured bilaterally distributed neural tissues (Fig. 7g-l). In accordance, we found that although *soxb* expression showed a fully radial pattern under the high-dose rhBMP4 treatment (supplemental Fig. S6m), a bilateral pattern was restored in a portion of embryos when the treatment was weaker (supplemental Fig. S6m’ and m’’).

## Discussion

While the DV patterning mechanism is considered conserved across bilaterian clades (12), investigations on several spiralian representatives have revealed different results (13-18). Meanwhile, in various spiralian phyla, DV patterning is deeply integrated into a highly specialized organizer-driven developmental mode (23, 26-29), with the underlying molecular mechanism largely unknown. Together, these two lines of evidence indicate that spiralian DV patterning is essential to explore the conservation and plasticity of animal DV patterning and to decipher the molecular network underlying the spiralian D-quadrant organizer. In addition, as a crucial innovation at the origin of bilaterians, DV patterning affects crucial aspects of the bilaterian body plan (54, 55) and would have contributed to the flourishing of bilaterians since the Cambrian period. Given that the spiralian body plan has been suggested to be informative to infer the origin of bilaterians (56, 57), one can expect that investigations on spiralian DV patterning would reveal evidence indicating how the bilaterian body plan transits from that of its ancestor, which does not possess a DV axis.

In the present study, we explored the DV patterning of the mollusk *L. goshimai*; the major findings are presented in Fig. 8. We found that under the regulation of the D-quadrant organizer, a *bmp2/4*-*chordin*-based molecular network determined the BMP signaling gradient in very early embryos (approximately 60-cell stage, Fig. 8a). This gradient regulated DV patterning since early embryonic development was radialized when it was eliminated due to knockdown of *bmp2/4* or *chordin* (Fig. 8b). Examinations of influenced embryos further revealed correlations between the BMP signaling gradient (of early embryos) and the localization of blastopore and the organization of the larval nervous system, indicating regulatory effects of BMP signaling on the two essential tissues (Fig. 8b). The re-emerged asymmetry additionally indicated undetermined factors regulating polarized development along the 3B-3D axis that were independent of *bmp2/4* or *chordin* (Fig. 8b).

**Fig. 8.**
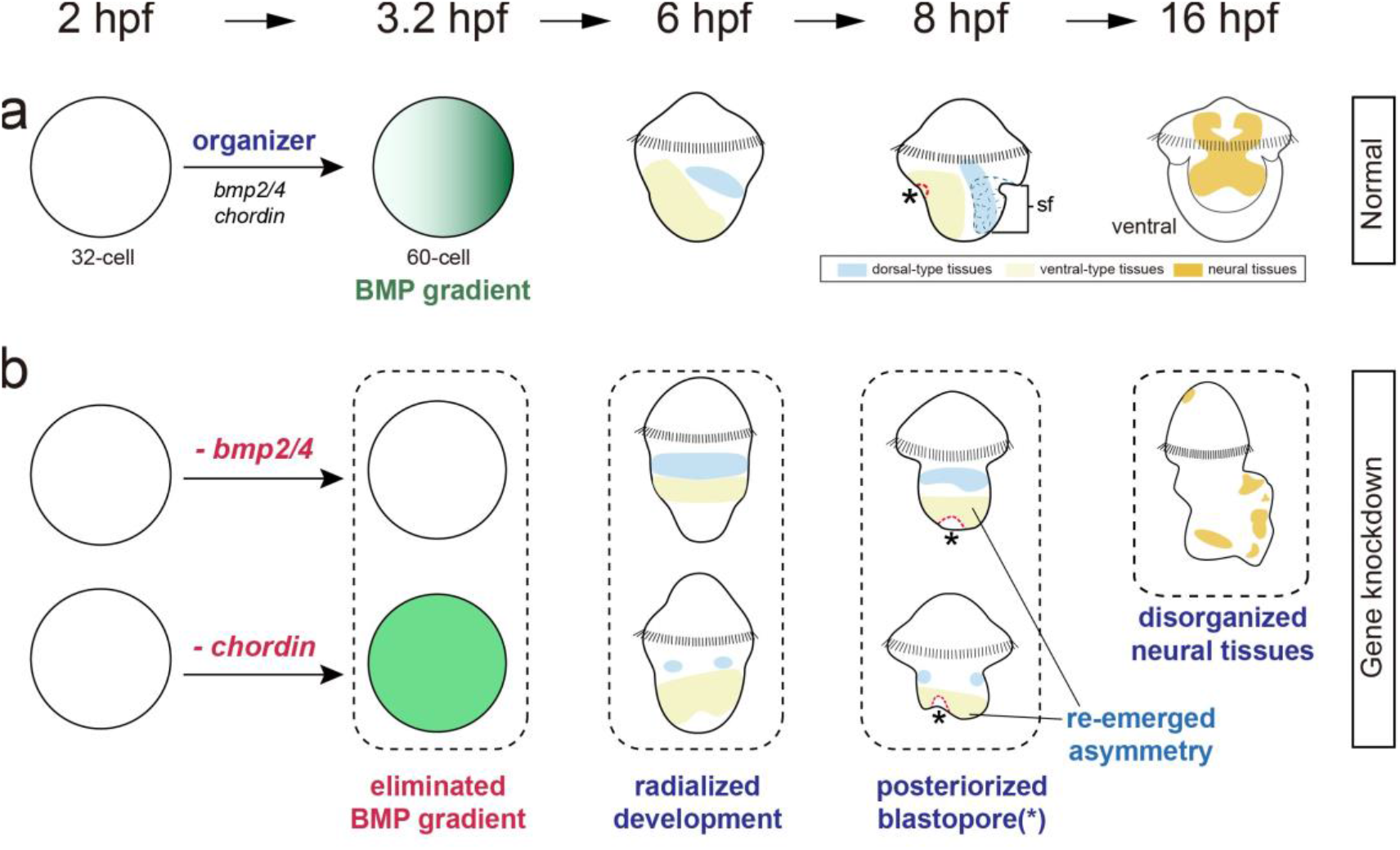
Schematic diagram showing the major findings of the present study. Focusing on *bmp2/4* and *chordin*, we revealed the roles of the two genes (i.e., BMP signaling) in organizer function (Figs. 2 and 3) and DV patterning of the mollusk *L. goshimai* (Fig. 5). A close look at the manipulated embryos revealed evidence showing the correlation between the BMP signaling gradient and the localization of blastopore (Fig. 6) and likely the organization of neural tissues (Fig. 7). Unexpected asymmetrical development re-emerged in the influenced embryos (Fig. 6). See more information in the text.

### L. goshimai DV patterning relies on bmp2/4 and chordin

A major issue in the studies of spiralian DV patterning is that researchers have reported very different results using different systems. Researchers concluded that spiralian DV patterning relies on *bmp2/4* (without known roles of *chordin*) (19, 20), other BMP ligand and antagonist (13), or even non-BMP signaling (14-18). Taken together, a paradox rises: while it is widely accepted that the common ancestor of bilaterians utilizes *bmp2/4* and *chordin* for DV patterning (12), this character is not revealed in the six spiralian representatives spanning three spiralian phyla (three annelids, two mollusks and one platyhelminth, Fig. 1a). This situation shows sharp contrast to the broad conservation of the DV patterning mechanism in other animal clades (1, 3, 5-7, 41). Here, we proved the roles of *bmp2/4* and *chordin* in DV patterning of the mollusk *L. goshimai* by showing that the knockdown of *bmp2/4* or *chordin* radialized early development (in 6-hpf embryos, Fig. 5). This result thus reveals a spiralian case retaining the conserved DV patterning mechanisms depending on *bmp2/4* and *chordin*, which is indicated but not revealed for a long time. Although this type of DV patterning mechanism has been revealed to be broadly conserved in most other animals, in the context of diverse reports on spiralian DV patterning, it conversely represents an unusual example. For other mollusks, a similar mechanism may be expected in the gastropod *Tritia* (20) and the bivalve *Crassostrea* (38, 39), given that current knowledge on their DV patterning mechanisms is consistent with a *bmp2/4-chordin*-based framework. Nevertheless, due to the very different experimental designs (see details below), we could not determine to what degree the differences between our conclusion and that in the gastropod *Crepidula* arguing no role of BMP signaling in DV patterning (14) would be explained by interspecies variations and the differences in experimental strategies.

After knockdown of DV patterning genes, dorsal/ventralization phenotypes are frequently observed in animals (e.g., *Drosophila*, vertebrates, hemichordates and echinoderms) (1, 3, 5, 41), including several spiralians (13, 19). However, *bmp2/4-* or *chordin*-knockdown phenotypes in *L. goshimai* showed relatively unusual characteristics. Despite the notably influenced cell specification, the major effect of gene knockdown is radialized development (Fig. 5), which we think is difficult to categorize as dorsal/ventralization. This result might be caused by technical reasons, e.g., the limited numbers of marker genes investigated. Indeed, a determined effect on tissue specification is suggested when taking *brachyury* expression into account. While some *brachyury* expression marked the ventral midline (Fig. 6r and s), we found that this part of *brachyury* expression was only retained in *chordin* knockdown embryos (Fig. 6t) but was undetectable when *bmp2/4* was inhibited (Fig. 6u). These results indicate the loss of the ventral midline in *bmp2/4*-knockdown embryos and could be interpreted to be a trend of dorsalization. However, this effect was not observed for the major dorsal and ventral-type tissues, namely, the shell field (*gata2/3*-positive tissues) and ventral plate (*soxb*-positive tissues) (Fig. 6a-q). In fact, even though the role of BMP signaling on ventral midline development was determined, this effect in *L. goshimai* is opposite to the situation in the spider *Achaearanea*, in which a comparable dorsalization phenotype is obtained when *chordin* (*sog*) is knocked down (6). This paradox, however, is easy to explain when considering that the “ventral” midline tissue of *L. goshimai* is actually formed on the dorsal side at the initial phases of its development and moved to its destined location later (supplemental Fig. S8c-e). Taken together, the largely uninfluenced cell specification and, in contrast, greatly disturbed tissue distributions in the influenced embryos (Figs. 5 and 6) suggest that the major role of BMP signaling in *L. goshimai* is to break down radial symmetry and regulate the originally AP-aligned tissues to move to dorsal and ventral sides, which together result in the formation of the DV axis. Given that DV patterning coincides with epibolic gastrulation in *L. goshimai*, we propose that the DV patterning signal may perform its role by regulating cell movement during gastrulation.

A close look at previous reports reveals that although not emphasized, radialized development without a certain trend of dorsal/ventralization was indeed observed in some animals. In the spider *Achaearanea, bmp2/4* (*dpp*) knockdown resulted in radialized development in which the genes that were either activated (e.g., *fkh*) or inhibited (e.g., *twist*) in normal ventral tissues all exhibited radial expression (6). In the cnidarian *Nematostella, bmp2/4* or *chordin* knockdown caused radial expression of some BMP components, and more importantly, both abolished the expression of the gene normally expressed on the opposite side of *chordin* expression (7). In these two cases, BMP signaling does not simply inhibit or promote the expression of particular genes. In contrast, polarized gene expression seems to be detectable only when a correct BMP signaling gradient is available. Collectively, our results as well as those previously reported suggest a role of BMP signaling in determining polarized distributions of gene expression without directly promoting or inhibiting the expression of the gene. In *L. goshimai*, this role seems to be achieved through regulating the movement of related cells; nevertheless, we do not suggest a similar process in *Achaearanea* or *Nematostella* given the insufficient details to reach a conclusion. Further investigations on additional species are necessary to explore the prevalence of this effect.

### Insights into the suggested diversity of spiralian DV patterning mechanisms

The very different reports in spiralian DV patterning undoubtedly suggest the diversity of the underlying mechanisms (13-20). However, it is notable that the experimental designs of these studies vary in many aspects: strategies to influence BMP signaling (MOs, small molecule inhibitors, exogenous BMP proteins), indicators of DV patterning (cell lineages, characteristic tissues, marker gene expression), and the developmental stages investigated (early embryos, larvae). Our results suggest that many of these factors would affect the results and may contribute to the differences in the conclusions.

One of our major findings is that the developmental stages investigated would be essential for the interpretations of the manipulated phenotypes. In *L. goshimai*, even though the developmental polarity was largely eliminated in early knockdown embryos (at 6 hpf, Fig. 5), asymmetrical development could re-emerge later (at 8 hpf, Fig. 6). In even later samples, some knockdown larvae exhibited a considerable degree of heterogeneity, and it was difficult to interpret whether and how DV patterning was influenced (e.g., at 15-16 hpf, shown Fig. 7). Such complicated phenotypes may be produced by the combined effects of influenced DV patterning, the amplification of disorganized early development (to varied degrees), etc. These results indicate that the effects on DV patterning may be underestimated if only a few or too late developmental stages were analyzed. Another difference among current studies is how researchers assess whether/how DV patterning is influenced. While cell lineage analysis has been particularly useful in studying spiralian development (23, 24, 26), the gene expression of particular blastomeres could be altered without detectable changes in cleavage patterns when influenced (e.g., *Tritia* Io2bRNA (20)). These data indicate that although cell lineage analysis is informative, gene expression data may be preferred when analyzing DV patterning. This strategy has also been widely used in DV patterning studies of many animals, including cnidarians, echinoderms and hemichordates (3, 7, 41).

The manners employed to influence BMP signaling would also be essential in studies on DV patterning. Given the extreme complexity of BMP signaling that includes multiple ligands and antagonists and is under tight temporal and spatial regulations (58, 59), manipulations on different nodes of the pathway could cause varied results. Indeed, we show that the phenotypes after *chordin* knockdown and rhBMP4 treatment somewhat differentiated from each other. Although they both caused enhanced BMP signaling and produced radialized development, only the *chordin*-knockdown embryos showed a posterior protrusion at 6 hpf (supplemental Figs. S5 and S6). Even radialized gene expression differed in the two groups: *chordin* knockdown caused circular *soxb* expression (Fig. 5n), and four symmetrical *soxb* expression was observed after treatment with 0.5 μg/ml rhBMP4 (supplemental Fig. S6m). These differences may be caused by nodes of the signaling that were manipulated. Theoretically, *chordin* knockdown causes one inhibited antagonist (several other uninfluenced) but the uninfluenced ligand repertoire, while rhBMP4 treatment results in one enhanced BMP ligand (still, several other uninfluenced) and the uninfluenced antagonist repertoire. Such differences in the nodes being manipulated may result in varied distributions and concentrations of BMP signaling, which would account for the differences between the resultant phenotypes. The complexity may further increase when considering that feedback effects exist between many nodes of the signaling pathway and that BMP signaling would function in multiple developmental processes (58, 59) (conversely, influenced DV patterning under both manipulations suggests the crucial role of *bmp2/4* and *chordin* in DV patterning of *L. goshimai*). Similarly, it is reasonable to assume that *bmp2/4* knockdown (specific inhibition of one BMP ligand) and the treatment of small molecular inhibitors (inhibition of all BMP ligands in general) would result in varied states of BMP signaling.

Taken together, given the many factors that may influence studies on spiralian DV patterning, we suggest analyzing marker gene expression at multiple developmental stages of representative species (while cell lineage analysis is still an important aspect). The results obtained though different manipulations should be compared to discriminate the effects caused by the interference of DV patterning and other biological processes. These efforts will help to make a better comparison among the current studies and understand the inconsistencies in their conclusions.

### DV patterning and the developmental mode in spiralians: organizer function and stereotype cleavage

A characteristic of spiralian DV patterning is that it depends on a D-quadrant organizer (26). The underlying molecular mechanisms of organizer function have received much attention (30-33, 37, 42, 60-62) but remain largely unknown. Although MAPK signaling is essential in organizer function (30, 31), the involved molecules are poorly understood. The demonstration of the involvement of *bmp2/4* in *Tritia* organizer function represents key progress toward an in-depth understanding of spiralian organizers (20). In the present study, we confirmed that *bmp2/4* played similar roles in *L. goshimai*, an equal cleaver, as its ortholog in the unequal cleaver *Tritia*. Although the manners of organizer activation and MAPK signaling dynamics would differ significantly between the two types of embryos (30, 31), the consistent employment of *bmp2/4* suggests a conserved molecular network underlying the organizer function of the two species. More importantly, we showed that after activation by the organizer, *bmp2/4* itself was not sufficient to establish the BMP signaling gradient in *L. goshimai* and that the formation of such a gradient required asymmetrically expressed *chordin*. This result supports our speculation of the indispensable role of *chordin* in organizer function. Our hypothesis regarding the regulatory relationships among the organizer, *bmp2/4* and *chordin* (Fig. 3) suggests that the canonical DV patterning molecular network has been deeply integrated into organizer function in spiralian development, thus consolidating the link between a highly clade-specific character (a D-quadrant organizer) and a conserved biological process (DV patterning) (20). This hypothesis can be tested in more spiralian lineages, which would be important to understand the unique developmental mode in spiralians (25, 26, 63). Nevertheless, we want to express a cautious attitude to infer the conservation/prevalence of this BMP2/4-based organizer function given that BMP signaling has been suggested to have no roles in DV patterning in two annelids and a mollusk (14, 15, 17, 18).

As a part of its inductive effects, the organizer regulates subsequent cleavage patterns (36, 61). Indeed, the different division manners of 3q blastomeres (larger 3a/b^2^ on 3B side versus larger 3c/d^1^ on 3D side) would be the first morphologically detectable polarity along the presumptive DV axis (36) (the 3B-3D polarity, supplemental Fig. S1d). This suggests that the highly stereotyped cleavage pattern may also participate in DV patterning. Intriguingly, although the BMP signaling gradient was eliminated when *bmp2/4* or *chordin* was knocked down, organizer formation and 3B-3D polarity in early embryos were not influenced (Fig. 2l and m). This provides an opportunity to explore the potential effects of this stereotype cleavage pattern. We detected asymmetry in late development of the manipulated *L. goshimai* embryos, which was along the 3B-3D axis in *chordin*-knockdown embryos (Fig. 5 & supplemental Fig. S5). Since 3B-3D polarity (supplemental Fig. S1d) would be the earliest and most evident polarity in the embryos, we propose that it may be related to late asymmetric development. This polarity may cause lineage-specific specifications (e.g., different fates of 3a/b^2^ and 3c/d^2^) or generate asymmetrical expression of other DV patterning genes on the 3B and 3D sides (e.g., *admp, tolloid*, etc.). Further investigations are required to clarify which factor is at work or whether the two factors act in combination. In any case, the asymmetrical development in the knockdown embryos indicates the roles of the organizer independent of *bmp2/4* or *chordin* and suggests the profound effects of the stereotype cleavage pattern on the development of *L. goshimai*.

### Evolution of the bilaterian body plan: the likely common signal regulating DV patterning, blastopore localization and neurogenesis

Bilaterians are phylogenetically close to cnidarians; it is suggested that the common ancestor of bilaterians exhibits a gastrula shape that shares many characters with extant cnidarians (55, 64). In particular, despite the tremendous variations, a generalized bilaterian possesses a DV axis, two digestive openings comprising a ventral mouth and a posterior anus, and a bilaterally organized nervous system. In contrast, their ancestor is suggested to lack a secondary axis and has a single anterior/posterior digestive opening and a radially organized nervous system. Obviously, the body plan of bilaterians experiences significant modifications compared to that of their ancestor. Several hypotheses suggest coordinated transitions of these structures during the early evolution of bilaterians, including innovation of the secondary axis, changes in digestive opening and formation of the directional, bilateral nervous system (54-57, 64). Nevertheless, it is unknown whether and how these transitions are coordinated at the molecular level. Our results show that these characteristics seem to be regulated by the same signal in *L. goshimai*, i.e., BMP signaling. We found that when signaling was influenced by knocking down *bmp2/4* or *chordin*, the embryos exhibited a body plan showing high similarities with the assumed bilaterian ancestor: no (or highly influenced) DV axis (radialized tissues distributed along the AP axis, Figs. 5 and 6), posteriorized blastopore (with the potential to develop to mouth, Fig. 6) and the lack of bilateral organization in the nervous system (likely in a radial pattern at early stages, as reflected by early *soxb* expression, Figs. 6 and 7). These results vaguely suggest that a common signal, manifested by the BMP signaling gradient at early embryonic stages, may contribute to the coordinated transitions of multiple key characters during the early evolution of bilaterians. In *L. goshimai*, such coordination seems to be achieved by a common role of BMP signaling in regulating cell movement and thus tissue distribution, although the details are to be elucidated. It would be intriguing to explore whether the coordinated development of these essential characteristics exists in other animal lineages, especially some spiralian and deuterostome lineages whose larvae possess ventral mouth and bilateral nervous system (e.g., other mollusks, brachiopods, echinoderms and hemichordates). From an evolutionary perspective, the cooption of these processes may have enhanced the fitness of the bilaterian ancestor and thus facilitated its success.

## Material and Methods

### Animals

Adults of *L. goshimai* Nakayama, Sasaki & Nakano, 2017, were collected from intertidal rocks in Qingdao, China. Spawning occurred after collection during the reproductive season (from June to August). During other seasons, algae were scraped from the surfaces of rocks inhabited by the limpets and cultured on plastic sheets under constant light. At 18-22 °C, the limpets fed these cultured algae could become sexually mature in several weeks. On some occasions, spawning was induced through elevated temperature, drying, rigorous water flow or sperm suspensions. The adult limpets were allowed to spawn in separate 100-mL cups, and the gametes were collected. Artificial fertilization was performed by mixing sperm and oocyte suspensions.

Fertilized eggs were incubated in filtered seawater (FSW) containing antibiotics (100 unit/mL benzylpenicillin and 200 μg/mL streptomycin sulfate) in an incubator at 25°C. The units of all developmental stages are in hpf except for the very early developmental stages (before the 64-cell stage). For *in situ* hybridization (ISH), samples at the desired developmental stages were fixed in 4% paraformaldehyde (1× PBS, 100 mM EDTA, and 0.1% Tween-20, pH 7.4), transferred to methanol and stored at -20°C until use. Older larvae (after 15 hpf) were anesthetized with 0.1% sodium azide or 125 mM magnesium chloride before fixation. Analyses of the samples were performed as previously described, including ISH (46), pSmad1/5/8 staining (41), phalloidin staining (65) and scanning electron microscopy (SEM) (39).

### Genes and MOs

*L. goshimai* gene sequences were first retrieved from a developmental transcriptome that we developed previously (46), and the orthologies were verified through subsequent phylogenetic analyses (supplemental Figs. 11-16). Translation-blocking MOs targeting *bmp2/4* (*bmp2/4* MO) and *chordin* (*chordin* MO1), as well as two negative control MOs (a muted *chordin* MO (control MO1) and a standard MO (control MO2)), were synthesized (supplemental text). In preliminary experiments, we confirmed that the two negative control MOs did not generate any detectable effects on the development of *L. goshimai* at the concentrations we used. Therefore, muted *chordin* MO (control MO1) was used as the negative control MO in most experiments. We also used another nonoverlapping MO to inhibit the *chordin* gene (*chordin* MO2, see the supplemental text) and confirmed that it generated a similar phenotype to that when using *chordin* MO1.

### MO microinjection

Microinjection was performed using a micromanipulator. The injection solutions contained 0.05% phenol red, 500 ng/μL FITC-conjugated dextran and 0.25 mM MO. No more than 1.5% of the oocyte volume of the injection solution was injected into the unfertilized oocytes (estimated by the diameter of the injected solution). After fertilization, successful injections were confirmed by the presence of green fluorescence inside the cells; embryos that exhibited no fluorescence were removed. In trials aiming to explore pSmad1/5/8 distribution, FITC-conjugated dextran was excluded from the injection solution to avoid causing relatively high background values in subsequent immunostaining. On these occasions, the injections were performed slowly and carefully to ensure that every injection was successful.

### Treatments with rhBMP4 or U0126

rhBMP4 (R&D Systems, USA; Cat. No. 314-BP) was resuspended in the suspending solution (0.2% BSA containing 4 mM HCl) at a concentration of 50 μg/mL and stored at -80°C according to the manufacturer’s instructions. Two doses of treatments were conducted, as determined by preliminary experiments testing a series of treatment parameters. Specifically, rhBMP4 was added at a final concentration of 0.5 μg/mL (the high-dose treatment) or 0.075 μg/mL (the low-dose treatment) immediately after fertilization, and the protein was eliminated from the culture system by three FSW washes at 6 hpf (supplemental Fig. S3). In the control groups, the same volume of suspending solution was added, and the same treatment time windows were used. The samples were collected at 6 hpf (for ISH) and 48 hpf (for investigations of eye development) before fixation.

U0126 (Beyotime, China; Cat. No. S1901) was dissolved in DMSO at a concentration of 25 mM and stored at -20°C. At the 16 -cell stage (approximately 1.7 hpf), the U0126 storage solution was added to seawater to a final concentration of 75 μM. In the control group, the same volume of DMSO was added. The embryos at the 60- to 64-cell stage (approximately 3.5 hpf) were transferred to FSW followed by three FSW washes to terminate the treatment and were then collected and fixed.

Oocytes from at least three females were used in every assay involving rhBMP4/U0126 treatment or MO injection (ISH, immunostaining, eye number investigation, and SEM), and we confirmed that maternal effects did not evidently influence the outcomes of most experiments. Limited maternal effects were observed in the low-dose rhBMP4 treatment group. In these groups, the broods derived from approximately 10% females developed two eye as those in normal development, contrasting with the broods derived from other females in which multiple-eye larvae were consistently observed. The eye development of these broods appeared to be less sensitive than that of other broods to low-dose *bmp2/4* treatment; they were not included in the subsequent analysis.

### Imaging

Images were recorded using a Nikon 80i microscope or an LSM 710 laser-scanning confocal microscopy system (ZEISS, Germany). The contrast and brightness of the images were adjusted using Photoshop software; when performed, such adjustments were applied to the whole image rather than to any particular regions.

## Acknowledgments

The authors are grateful to Dian-Han Kuo for engaging in thoughtful discussions throughout the course of the study and for critically revising the draft. The authors thank Menglu Cui for performing some ISH experiments, Bo Dong and Grigory Genikhovich for their discussions and comments on the draft, Ferdinand Marlétaz for providing the assembled transcriptomic data of various spiralians and for the helpful discussions, and Patrick Müller, Ulrich Technau and Lingyu Wang for their helpful discussions. This study was funded by the National Key R&D Program of China (2018YFD0900104) grant to P.H., the Marine S&T Fund of Shandong Province for Pilot National Laboratory for Marine Science and Technology (Qingdao) (2018SDKJ0302-1) grant to B.L., the China Agriculture Research System (CARS-49) grant to B.L., the National Natural Science Foundation of China (NO. 42076123 and 31472265 grant to B.L. and NO. 41776157 grant to P.H.) and the Youth Innovation Promotion Association CAS (2018239) grant to P.H.

