## Supplementary materials for "Molluscan dorsal-ventral patterning relying on *bmp2/4* and *chordin* provides insights into spiralian development and bilaterian evolution"

**Supplementary materials:**

- **Supplemental Text**
- **Supplemental Figures S1-S16.**
- **Caption for supplemental dataset 1**
- **Supplemental References**

### Supplemental Text

***MO oligos***

The MO oligos were designed by Gene Tool, LLC (https://www.gene-tools.com/). The MOs that we used in the present study are below.

(1) *bmp2/4* MO (for *bmp2/4* knockdown): AATCTGCGATCATAGTGGACACAAC

(2) *chordin* MO1 (for *chordin* knockdown):
AACTGCTAAATCCATAGGAGACATG

(3) *chordin* MO2 (for *chordin* knockdown):
TGATCGCCCAGTATGGGAAATTACC

(4) Control MO1 (negative control): AAgTGgTAAATCgATAGcAcACATG (this MO is derived from *chordin* MO1, with six residues muted [indicated by lowercase letters]. This MO is predicted to lack the ability to bind to *chordin* mRNA.)

(5) Control MO2 (negative control, Standard Control MO):
CCTCTTACCTCAGTTACAATTTATA

*bmp2/4* MO is designed to block translation of *bmp2/4*; the location of the MO-binding site is shown below (the start codon is shown in bolded letters):

```
                          bmp2/4 MO
MO              3’ CAACACAGGTGATACTAGCGTCTAA 5’
                   |||||||||||||||||||||||||
bmp2/4 mRNA  ...GGTGTTGTGTCCACTATGATCGCAGATTTTTA...
```

*chordin* MO1 and *chordin* MO2 are designed to block translation of *chordin*; the locations of the MO-binding sites are shown below (the start codon is shown in bolded letters):

```
                         chordin MO2                            chordin MO1
MOs           3’ CCATTAAAGGGTATGACCCGCTAGT 5’           3’ GTACAGAGGATACCTAAATCGTCAA 5’
                    |||||||||||||||||||||||||                 |||||||||||||||||||||||||
chordin mRNA ...GTGGTAATTTCCCATACTGGGCGATCACCCAGCCCGCTGTTTAAACATGTCTCCTATGGATTTAGCAGTTTGT...
```

The results we show in the figures were generated using *chordin* MO1, *bmp2/4* MO and Control MO1 given that the two *chordin* MOs generated similar phenotypes and the two control MOs did not show detectable effects on the development of *L. goshimai* at the concentrations we used.

***Determination of the identity of the organizer in L. goshimai***

Although the organizer of many spiralians can be readily identified by activated MAPK signaling (reflected by the immunostaining of double phosphorylated ERK (dpERK)), we failed to label dpERK in *L. goshimai* embryos due to technical problems. No signals were produced, although we tried all published protocols we could retrieve. Nevertheless, we could generally determine the organizer of *L. goshimai* to be 3D based on the following characteristics:

(1) The cleavage patterns (from 1 to 64 cells) were almost identical to *Patella* (1): 1-cell (zygote) - 2-cell (AB + CD) - 4-cell (A + B + C + D) - 8-cell (1q + 1Q) - 16-cell ($1q^{1}$ + $1q^{2}$ + 2q + 2Q) - 32-cell ($1q^{11}$ + $1q^{12}$ + $1q^{21}$ + $1q^{22}$ + $2q^{1}$ + $2q^{2}$ + 3q + 3Q) - 40-cell (division of $1q^{21}$ and $1q^{22}$) - 52-cell (division of $1q^{11}$, $1q^{12}$ and $2q^{1}$) - 60-cell (division of $2q^{2}$ and 3q) - 63-cell (division of 3Q except 3D) - 64-cell (division of 3D).

(2) A prolonged interdivision phase after the fifth cleavage (from 32- to 40-cell stage; which took approximately 30 minutes compared with less than 20 minutes in previous stages).

(3) Asymmetrical cellular arrangements after the formation of the presumed organizer: a more centered macromere (the assumed 3D blastomere) compared to the other three (3A-3C) since the 52-cell stage and a characteristic four-cell arrangement pattern at the vegetal pole at the 60-cell stage ($2d^{22}$, $3c^{2}$, $3d^{2}$ and 3D; see supplemental Fig. S1a and c).

(4) When inhibiting MAPK signaling using U0126 from the 16-cell to 64-cell stage (as conducted in many mollusks; see the Materials and Methods section), the

asymmetrical characteristics at the cellular level mentioned in (3) could be eliminated, and a radial distribution was produced (Fig. 2a and Supplemental Fig. S3).

***Primers used in WMISH***

Specific primers containing the T7 promoter sequence (taatacgactcactataggg) in their 5′-upstream regions were used to generate complementary DNA (cDNA) fragments of genes. Sequences of the primers are shown below.

***bmp2/4,*** F: 5'ACCAGAAGCAAATTCCTCAAGT3';
R: 5'CCCTCTACTACCATATCTTGATAG3';
***chordin,*** F: 5'AGCCAGTTCCAAGACCAGATTCA3';
R: 5'GAACCACAGCAGAACCTTCATCAA3';
***brachyury,*** F: 5'TACGAAGAACGGCAGGAGAATGT3';
R: 5'TGAGACACAACACTAGAGGCTGAG3';
***foxa,*** F: 5'GCTTATCACAATGGCTATTCAACAGTC3';
R: 5'CGTGGAATGAGGAGCGTAGGT3';
***gata2/3,*** F: 5'CCATCCGAAGACATAAGTGACA3';
R: 5'ATGATAGAGACCACAGGCGTTA3';
***soxb,*** F: 5'TTGCGACCGATCCTAGAGTGA3';
R: 5'TCCGTCTGTGGTGGCTTGT3';
***elav,*** F: 5'ACCACAGACAATGACACAAGATGAT3;'
R: 5'ACTCGGTTGCCTAATGTGAATCC3';
***pax6,*** F: 5'GGTAGTGCTCAGGCGTGTAGT3';
R: 5'GCTCCATTGACAACTTCTCTTCTCT3'.

### Supplemental figures

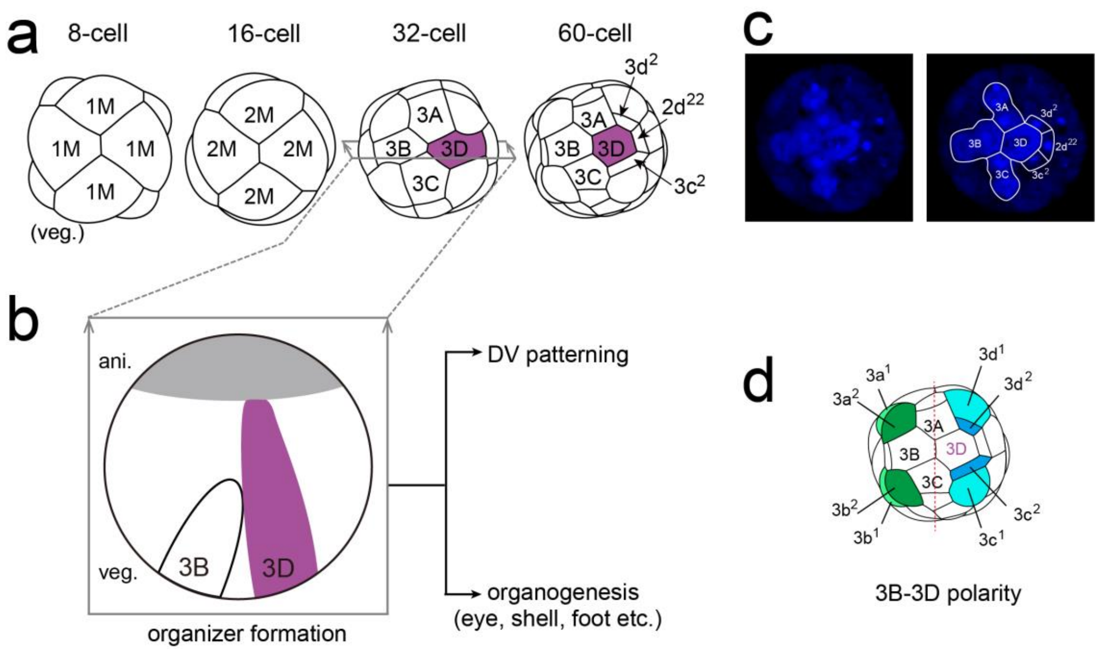


**Fig. S1 *L. goshimai* shows highly conserved spiral cleavages involving a D-quadrant organizer. a.** Schematic diagrams showing the early cleavages of *L. goshimai* from the 8- to 60-cell stage (vegetal views). All macromeres located at the vegetal pole are functionally equivalent initially (1M and 2M at the 8- and 16-cell stages). A macromere is induced to form the organizer at the 32-cell stage; it is named 3D, and the other macromeres are designated 3A-3C. At the 60-cell stage, due to the inductive effects of the organizer, a characteristic 4-cell arrangement (3D, $2d^{22}$, $3c^2$ and $3d^2$) is discriminable at the vegetal pole. **b**. In equal-cleaving species, such as *L. goshimai*, the formation of the organizer (3D) is induced through contacts with the micromeres and is marked by MAPK signaling (purple). The organizer then induces two processes that largely overlap: DV patterning and organogenesis. **c**. Early embryo of *L. goshimai* at the 60-to-63-cell stage showing the characteristic 4-cell arrangement at the vegetal pole. In this sample, the 3A-3C blastomeres are in the course of division. **d**. The asymmetry on 3B and 3D sides at the cellular level recognizable in early embryos (3B-3D polarity). This polarity is mainly reflected by the 3q blastomeres (highlighted by green and blue colors), in addition to the difference between 3B and 3D blastomeres.

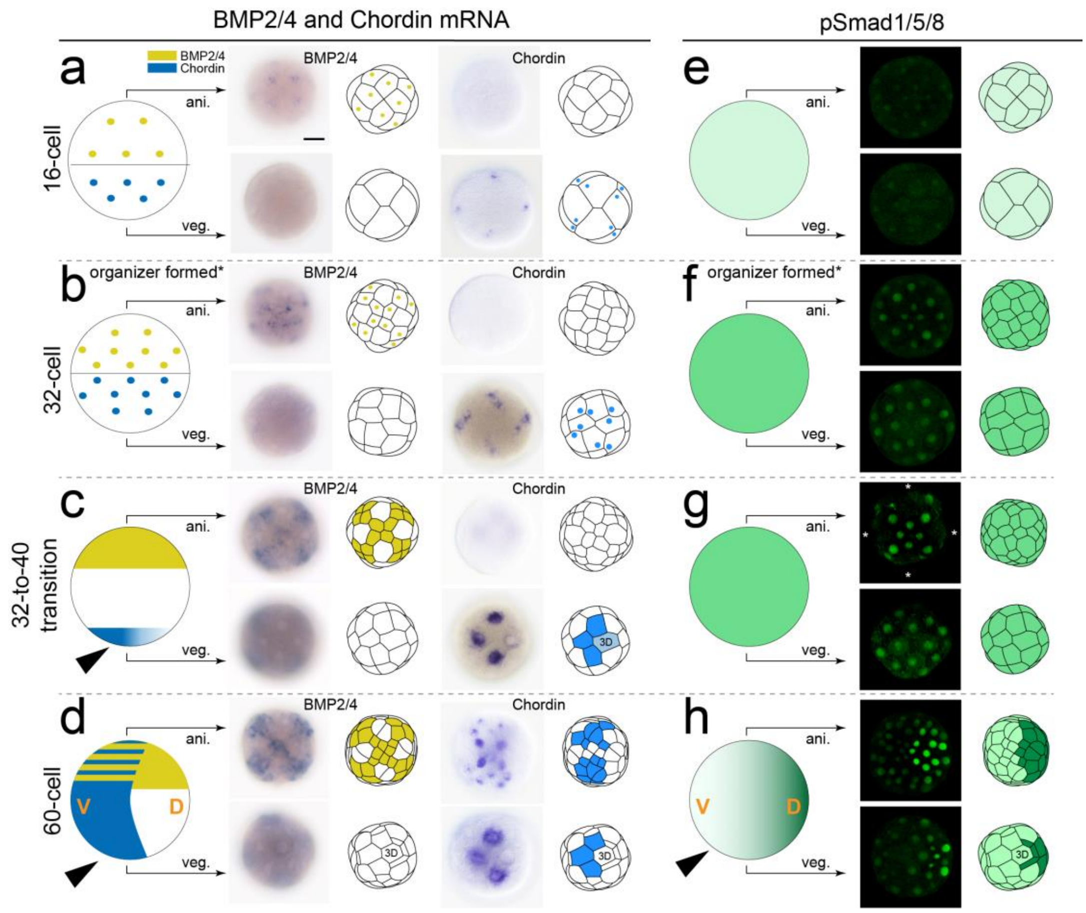


**Fig. S2 Dynamic changes in *bmp2/4* and *chordin* mRNA expression (a-d) and BMP signaling (e-h) during the periods before and after organizer formation.** In each panel, a diagram showing the overall state of BMP signaling or gene expression is presented on the left side (yellow for *bmp2/4* mRNA and blue for *chordin* mRNA in **a-d**, green for BMP signaling in **e-h**). Note that all schematic diagrams are lateral views. On the right side of the panel, the animal (ani.) and vegetal (veg.) views of the stained embryo are shown along with corresponding diagrams. The state of BMP signaling is reflected by pSmad1/5/8 signals (confocal projections). The organizer is distinguishable based on the characteristic 4-cell arrangement at the vegetal pole at the 60-cell stage (**d** and **h**), while the organizer at an earlier stage (in **c)** is distinguished based on decreased *chordin* expression in the blastomere. Organizer formation at the 32-cell stage is indicated by a prolonged pause in cell division after the fifth cleavage (asterisks in **b** and **f**, see supplemental text). Essential events that break down radial symmetry are highlighted by arrowheads, including changes in both *chordin* mRNA expression (**c**, **d**) and BMP signaling (**h**). At the 60-cell stage, the BMP signaling gradient (**h**) is complementary to the *chordin* mRNA expression pattern (**d**), which establishes a molecular DV axis. In panel **g**, the primary trochoblasts (white asterisks) lacked pSmad1/5/8 signals since the cells were dividing; we found that pSmad1/5/8 staining became detectable when cell division was complete at the 40-cell stage.

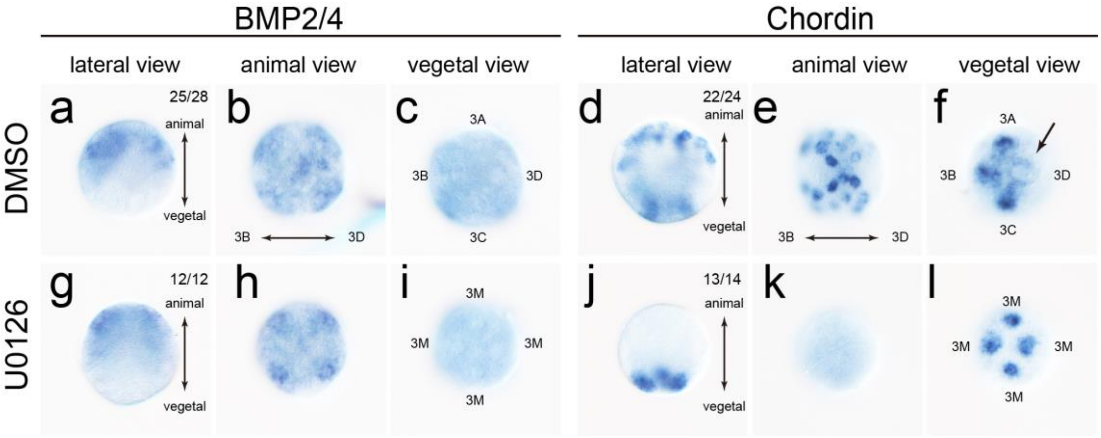


**Fig. S3 Changes in *bmp2/4* and *chordin* mRNA expression after U0126 treatment.** The samples are at the 60- or 63-cell stage. When organizer formation was inhibited by U0126 treatment (Fig. 3j), *bmp2/4* expression did not change in general (**a-c**, **g-i**). In contrast, significant changes in *chordin* expression occurred (**j-l**), which failed to transit into an asymmetrical pattern, as in normal embryos (**d-f**). In particular, expression in the animal pole was not detected (**k**, compare to **e**), and expression in the vegetal pole showed a radial pattern (**l,** compare to **f**), assembling the pattern in earlier stages (Fig. S4b). This result reveals that *chordin* is regulated by the organizer. However, since *chordin* expression was not eliminated in **j-l**, it seems that such regulation is restricted to generating an asymmetrical pattern but not maintaining basic chordin expression. Another notable fact is that the radial-to-asymmetrical transition of *chordin* expression included not only the downregulation of the expression in the organizer (arrow in **f**) but also the activation of the gene in some animal blastomeres (**e**). Regulation from the organizer to animal blastomeres, if it exists, may be explained by the relatively large volume of the organizer that may be spatially adjacent to animal blastomeres inside the embryo. Nevertheless, we do not deny other potential regulators that regulate *chordin* expression in animal blastomeres.

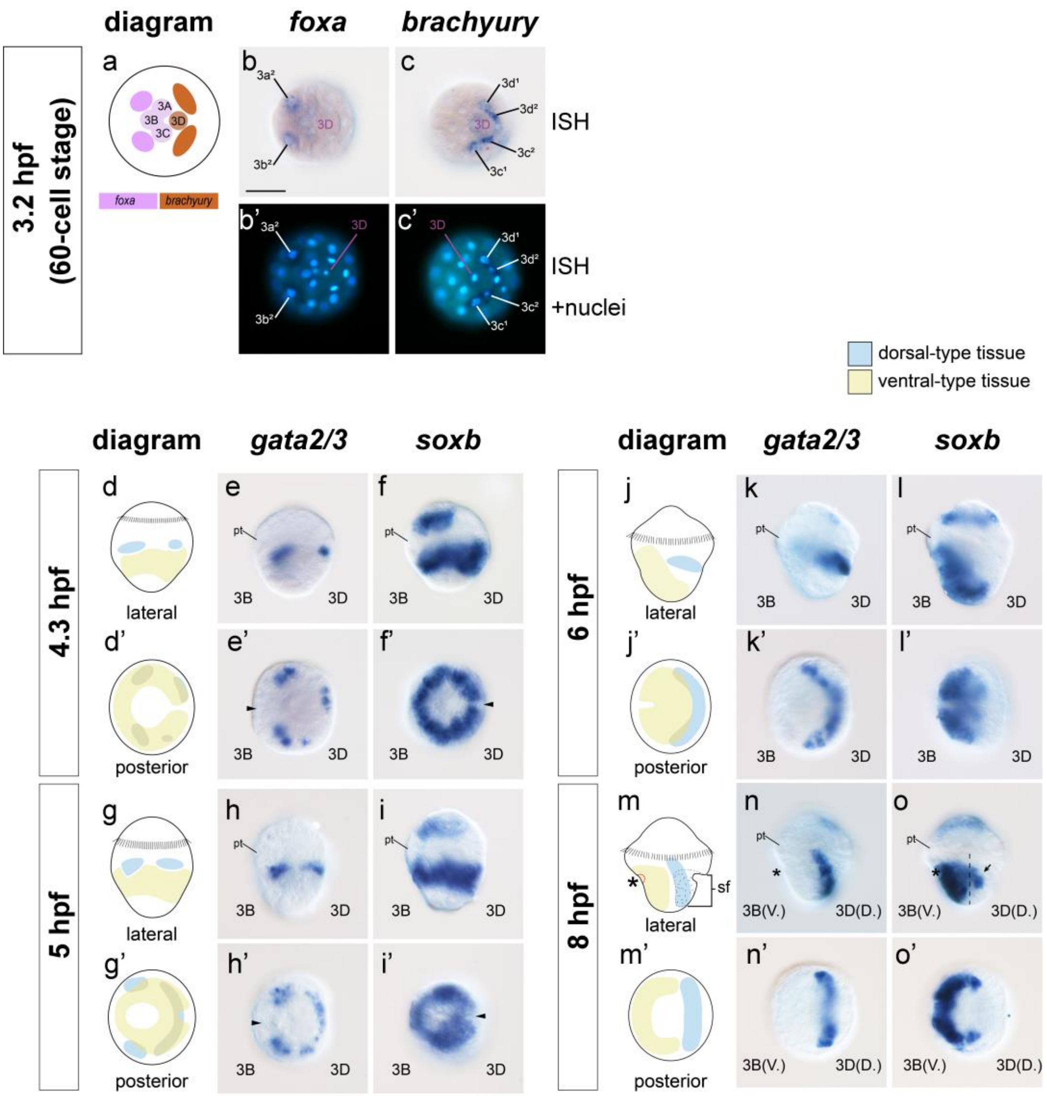


**Fig. S4 DV patterning of *L. goshimai* based on analysis of marker gene expression.** At the very beginning of DV patterning (60-cell stage), asymmetrical expression of *foxa* and *brachyury* could be detected (**a-c**). Panels **d-o** show the expression of *gata2/3* and *soxb*, markers of the shell field and ventral plate, respectively, thereby reflecting the normal DV patterning process. The development of the two tissues exhibits two major characteristics. First, they are initially aligned along the anterior-posterior axis (**d-f**, **g-i**) and gradually move to their destined locations on the dorsal and ventral sides (**j-l**, **m-o**). Second, they show largely circular organizations at early developmental stages (**d'-f'**, **g'-i'**). However, dorsal-ventral asymmetry could still be detected, which is reflected by the expression gaps along the presumptive DV axis (arrowheads in **e'-f'** and **h'-i'**). The arrow in **o** indicates a part of *soxb* expression that was likely distributed in dorsal-type tissues (see also in ref (2)). The asterisks in **m-o** indicate the blastopore. sf, shell field.

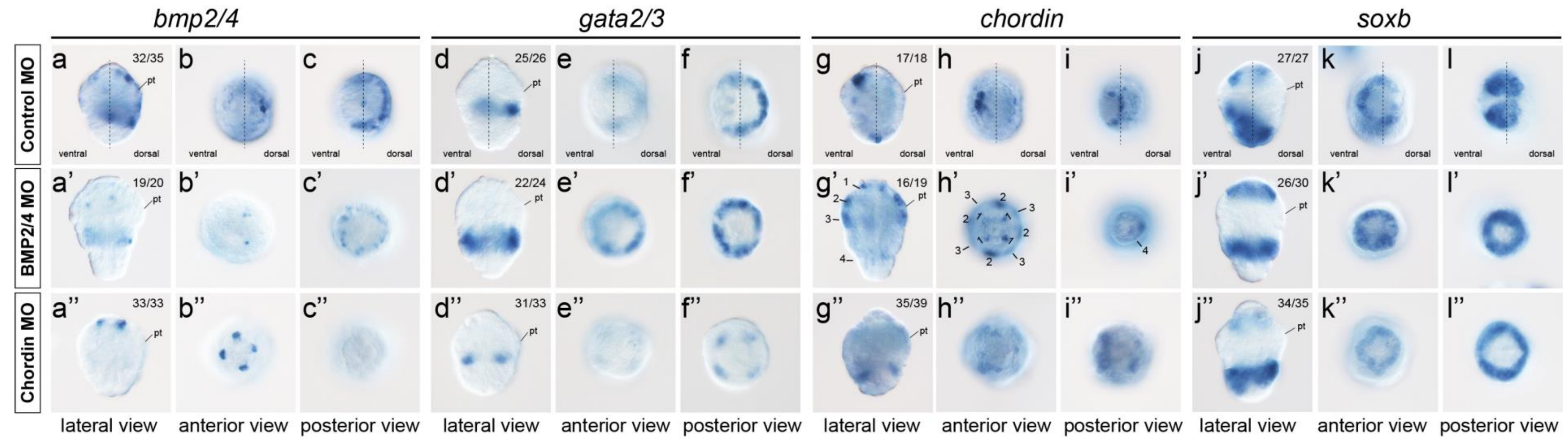


**Fig. S5 6-hpf embryos show radial development after *bmp2/4* or *chordin* knockdown.** All panels show 6-hpf gastrulae. At this stage, the four genes investigated showed evident asymmetrical expression in the control group. In contrast, the genes exhibit general radial expression patterns after gene knockdown (when detectable). The numbers (1-4) in **g'-i'** indicate *chordin* expression in four "tiers" of cells.

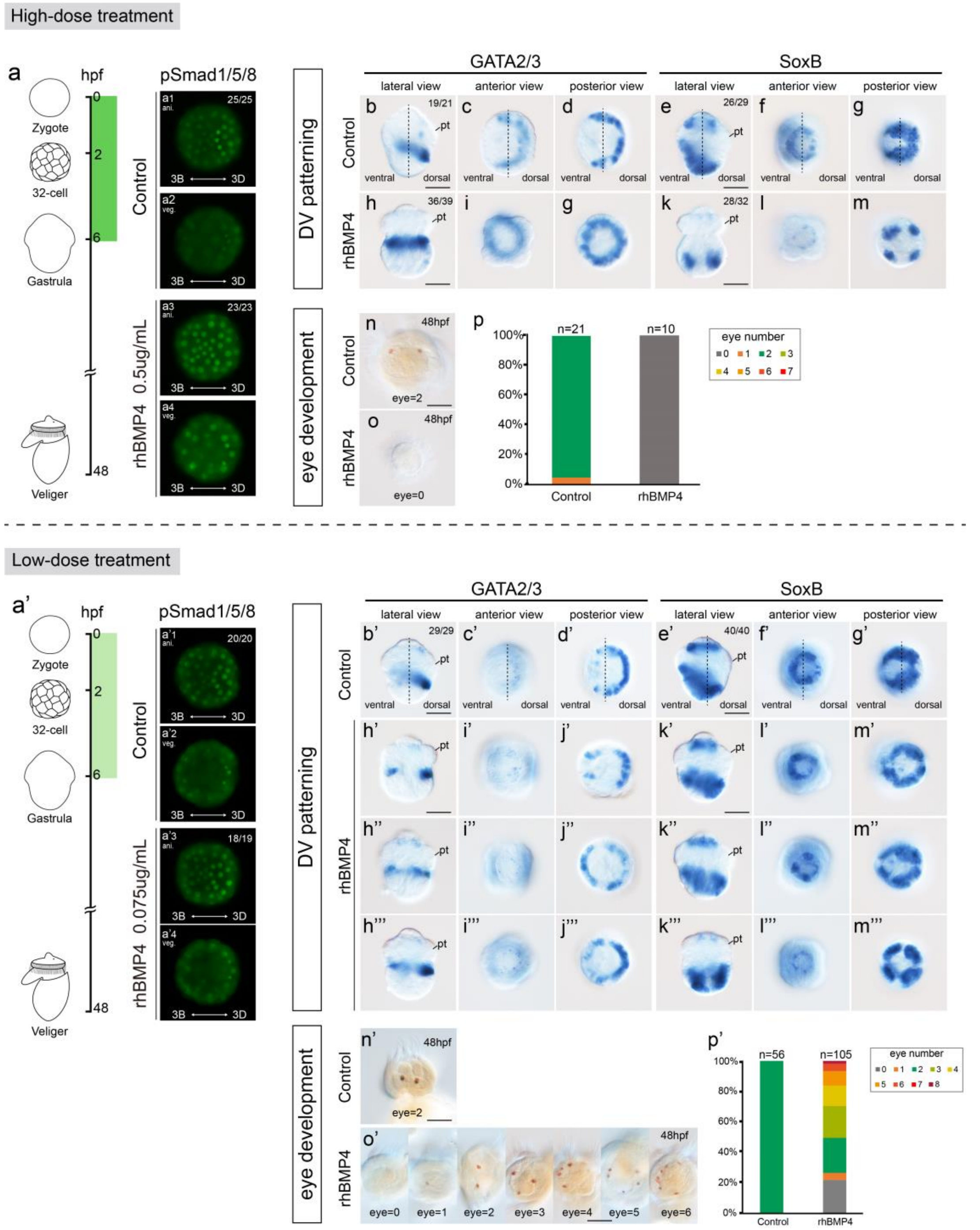


**Fig. S6 Phenotypes after recombinant human BMP4 (rhBMP4) treatment.** Two different doses of rhBMP4 treatment were conducted (**a** and **a'**). Altered BMP signaling, DV patterning and larval eye numbers were investigated after the treatments. Under high-dose treatment (**a**), BMP signaling was universally distributed in the whole embryo, reflected by the universally distributed pSmad1/5/8 (**a3-a4**). The radial expression of marker genes *gata2/3* and *soxb* (**b-m**) indicates disrupted DV patterning; no larval eyes were developed (**n-p**). Under low-dose treatment (**a'**), BMP signaling was enhanced, but polarity along the 3B-3D axis was still detectable (**a3'-a4'**).

Although DV patterning was disrupted in some embryos (**b’”-m’”**), the radial expression of marker genes was broken down in others (moderately in **b”-m”** and severely **in b’-m’**). The most different result compared to the high-dose group was that extra eyes were observed after low-dose treatment (**n’-p’**), revealing dose-dependent effects of BMP signaling on larval eye development.

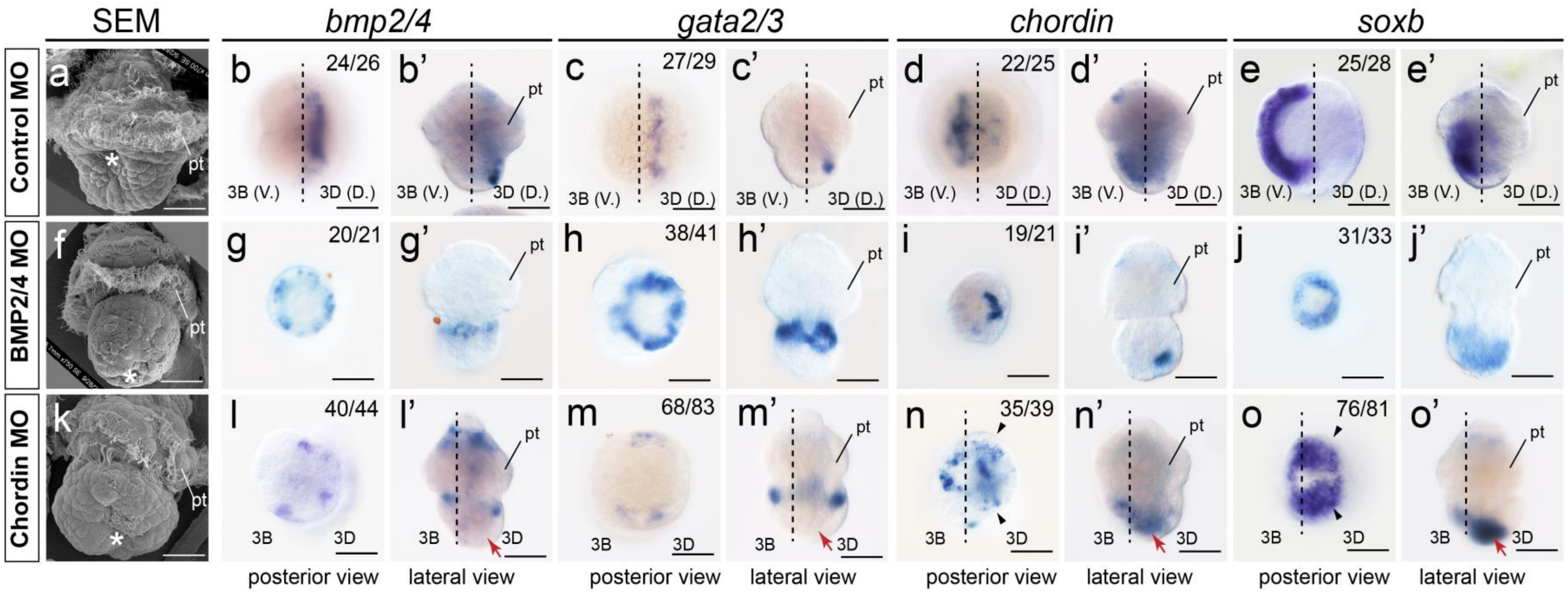


**Fig. S7 8-hpf embryos after *bmp2/4* or *chordin* knockdown.** All panels show 8-hpf embryo. SEM revealed that a common characteristic of knockdown embryos is that the blastopore is posteriorized (asterisks in **f** and **k**), contrasting with the ventral blastopore in the normal embryo (**a**). The four genes investigated show almost exclusively dorsal or ventral expression in the control group (**b-e**). The embryos injected with *bmp2/4* MO generally showed radial development (**g-j**) at an earlier stage (Fig. 5) despite the polarity of *chordin* expression (**i**). In contrast, the embryos with *chordin* knockdown showed asymmetrical development along the 3B-3D direction (**l-o**), which was evidently reflected by the expression of *chordin* (**n, n'**) and *soxb* (**o, o'**). Moreover, the posterior part of the embryo is further divided into two bilateral lobes (arrowheads in **l-o**). pt, prototroch. The bars represent 50 μm.

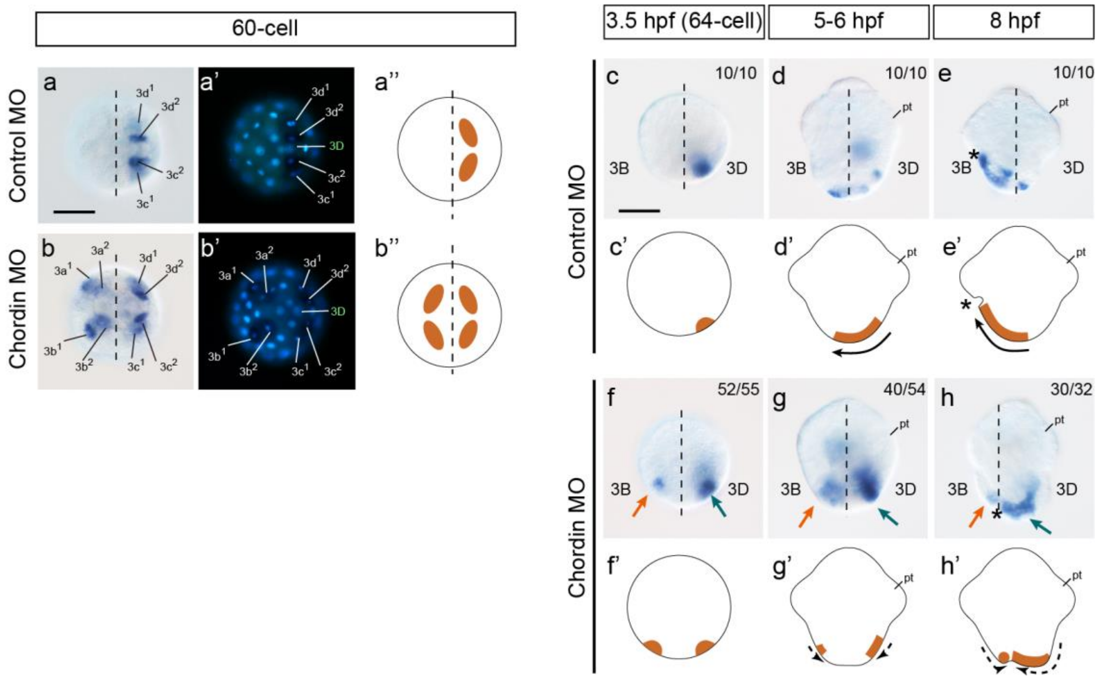


**Fig. S8 *brachyury* expression in normal embryos and those after *chordin* knockdown.** At the 60-cell stage (**a-a'** and **b-b'**, vegetal views), vegetal blastomeres are recognizable due to the characteristic 4-cell arrangement. While *brachyury* expression is detectable only on the 3D side in a normal embryo (**a-a'**), additional expression in a mirror manner is observed on the opposite side (3B side) with *chordin* knockdown (**b-b'**). The changes in gene expression mainly occurred in 3q blastomeres (**a'** and **b'**); however, no evident morphological differences were observed between normal and knockdown embryos. In subsequent developmental stages, extensive cell movements occur, marking a characteristic of epibolic gastrulation (arrows in **c'-e'**). After *chordin* knockdown, *brachyury* expression (**f-h**) suggests duplicated epiboly (dashed arrows in **f'-h'**); nevertheless, this speculated cell movement mode requires further investigation for confirmation. More importantly, during this period (**g**, **h**), *brachyury* expression on the 3B side (red arrows) was downregulated, while that on the 3D side expanded significantly (green arrows). These results help to determine the orientation of the embryos after *chordin* knockdown.

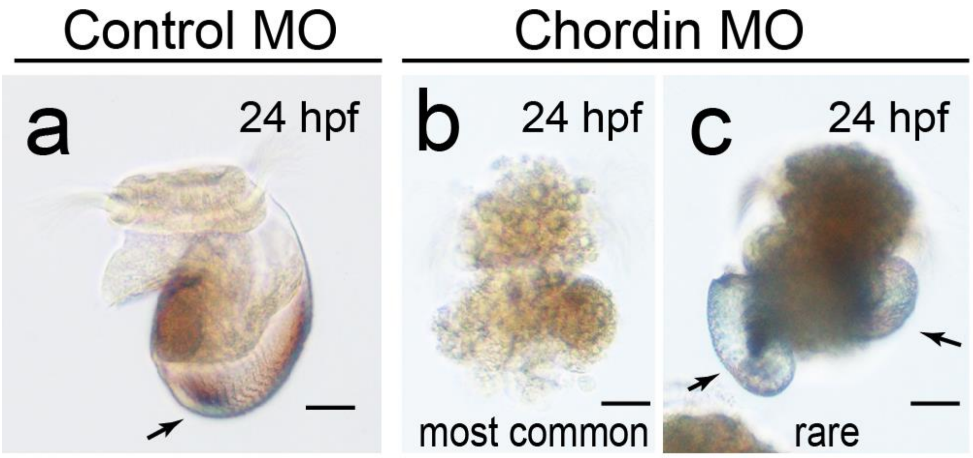


**Fig. S9 Duplicated larval shells were developed in rare cases after *chordin* knockdown.** Panel **a** shows a normal veliger larva at 24 hpf. Most of the *chordin* knockdown embryos did not develop discriminable posttrochal organs (**b**), except that a shell was developed occasionally. However, in a brood among over hundreds of replicates that we performed, duplicated shells were developed in two of 41 recorded larvae, each shell of which even showed a spiral shape (highlighted by arrows in **c**). This result indicates that the pseudotwin phenotype has the potential to develop duplicated structures (at least duplicated shell fields). The development of duplicated shells, however, never reemerged when we repeated the experiments using varied MO concentrations or a different MO, indicating a very special genetic background of that brood. Nevertheless, despite the poor reproducibility, we think it is necessary to describe this duplicate-shell phenotype (probably the first report of gastropod larvae with duplicated shells) since it may provide insights into DV patterning, shell formation and their relationships.

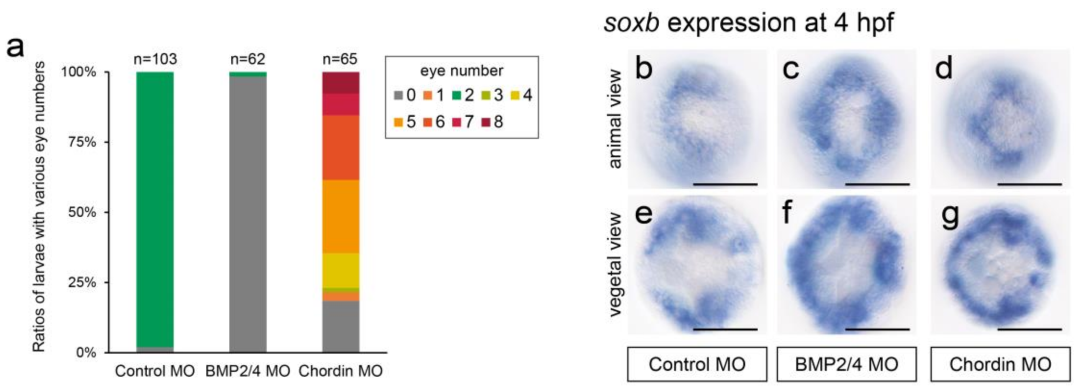


**Fig. S10 Effects of BMP signaling on eye development (a) and early *soxb* expression (b-g).** The bars represent 50 µm.

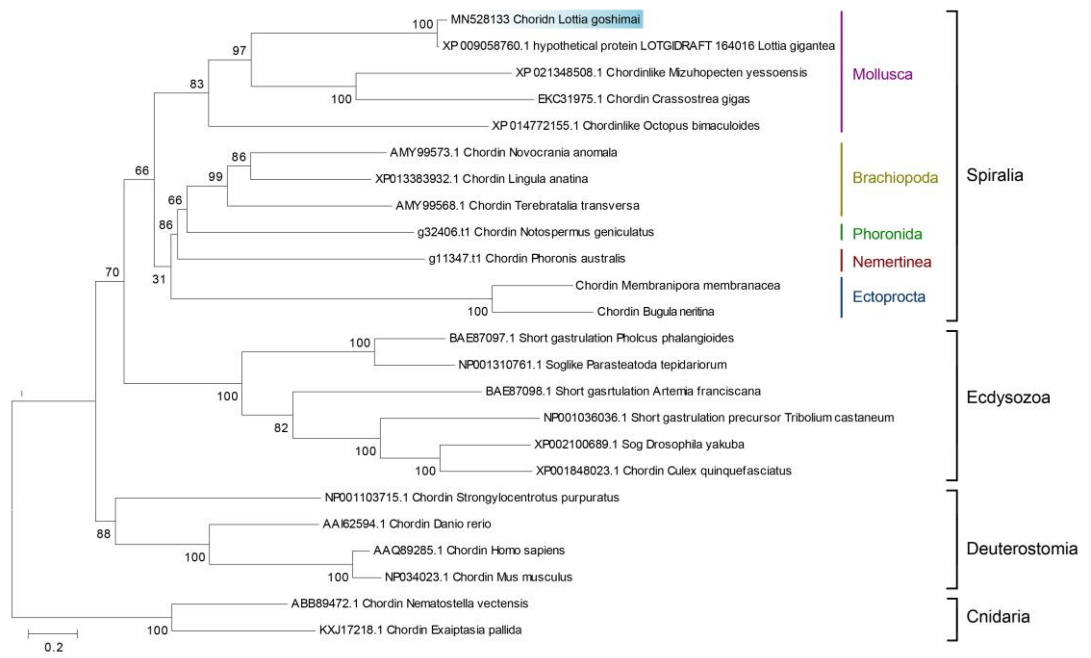


**Fig. S11 Maximum likelihood (ML) tree of *chordin* genes.** The sequences were retrieved from representative animals with special attention to spiralians. Phylogenetic analysis was performed using MEGA 6.0 with the whole amino acid sequences. The WAG + G + I model was estimated to be the best-fitting evolutionary model and was thus used in the analysis. The numbers at the nodes represent the bootstrap percentages from 1000 replicates. All sequences used in this phylogenetic analysis are provided in supplemental dataset 1.

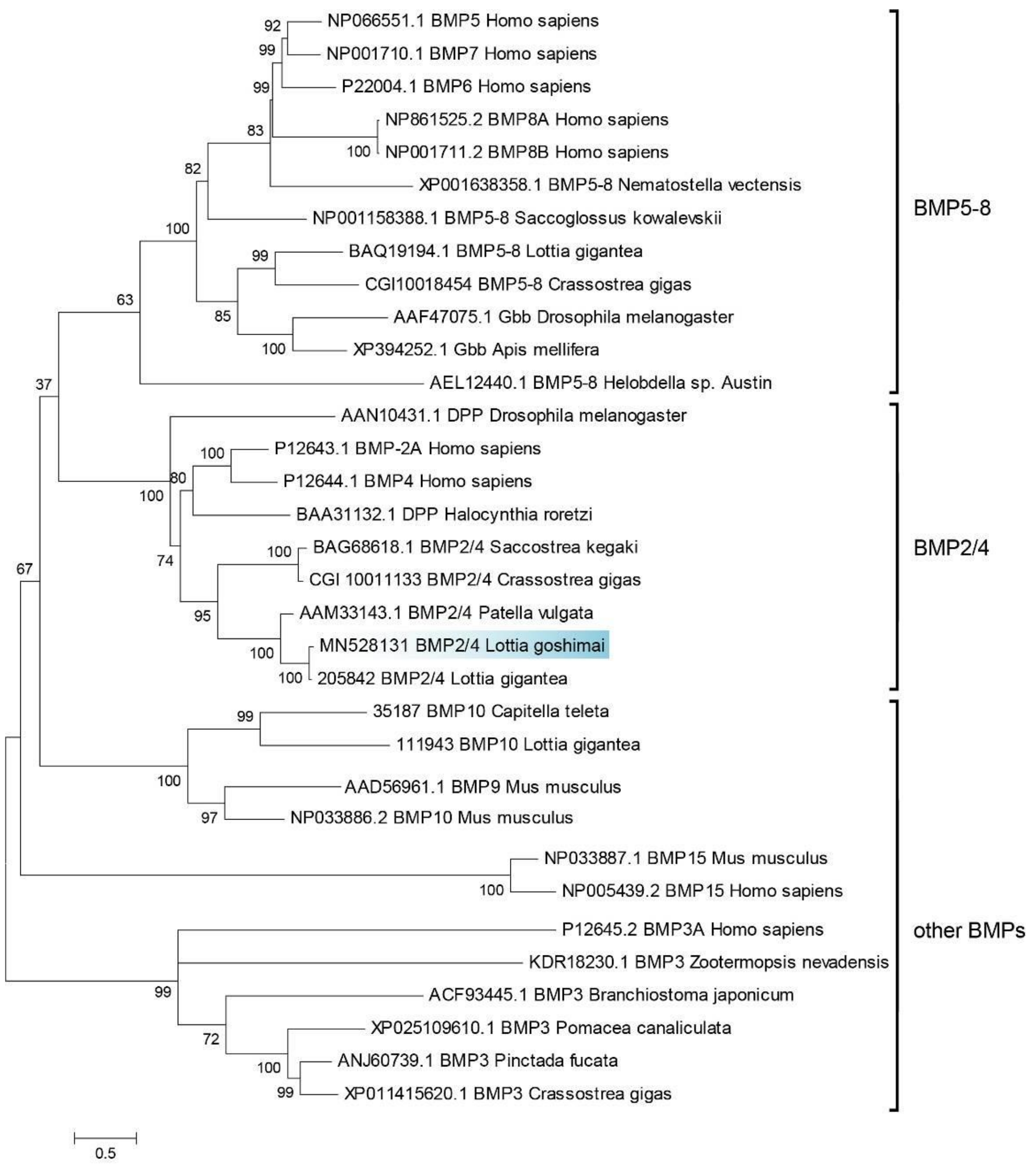


**Fig. S12 ML tree of BMP genes.** Phylogenetic analysis was performed using MEGA 6.0 with the whole amino acid sequences. The WAG + G + I + F model was estimated to be the best-fitting evolutionary model and was thus used in the analysis. The numbers at the nodes represent the bootstrap percentages from 1000 replicates.

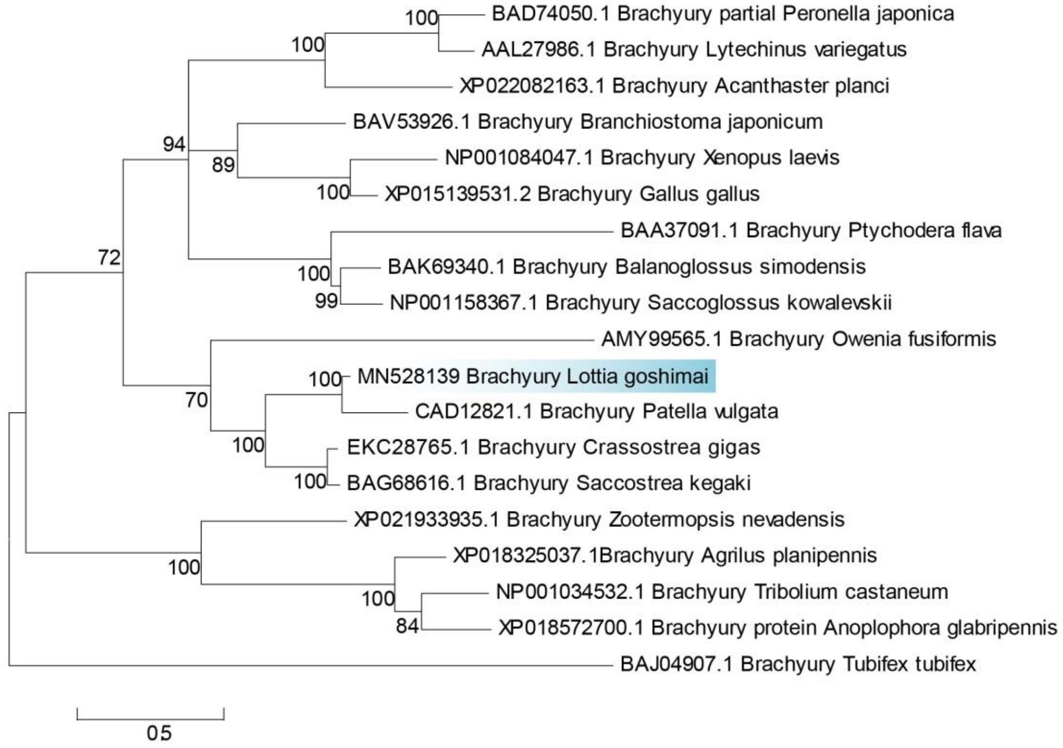


**Fig. S13 ML tree of *brachyury* genes.** Phylogenetic analysis was performed using MEGA 6.0 with the whole amino acid sequences. The JTT + G + I model was estimated to be the best-fitting evolutionary model and was thus used in the analysis. The numbers at the nodes represent the bootstrap percentages from 1000 replicates.

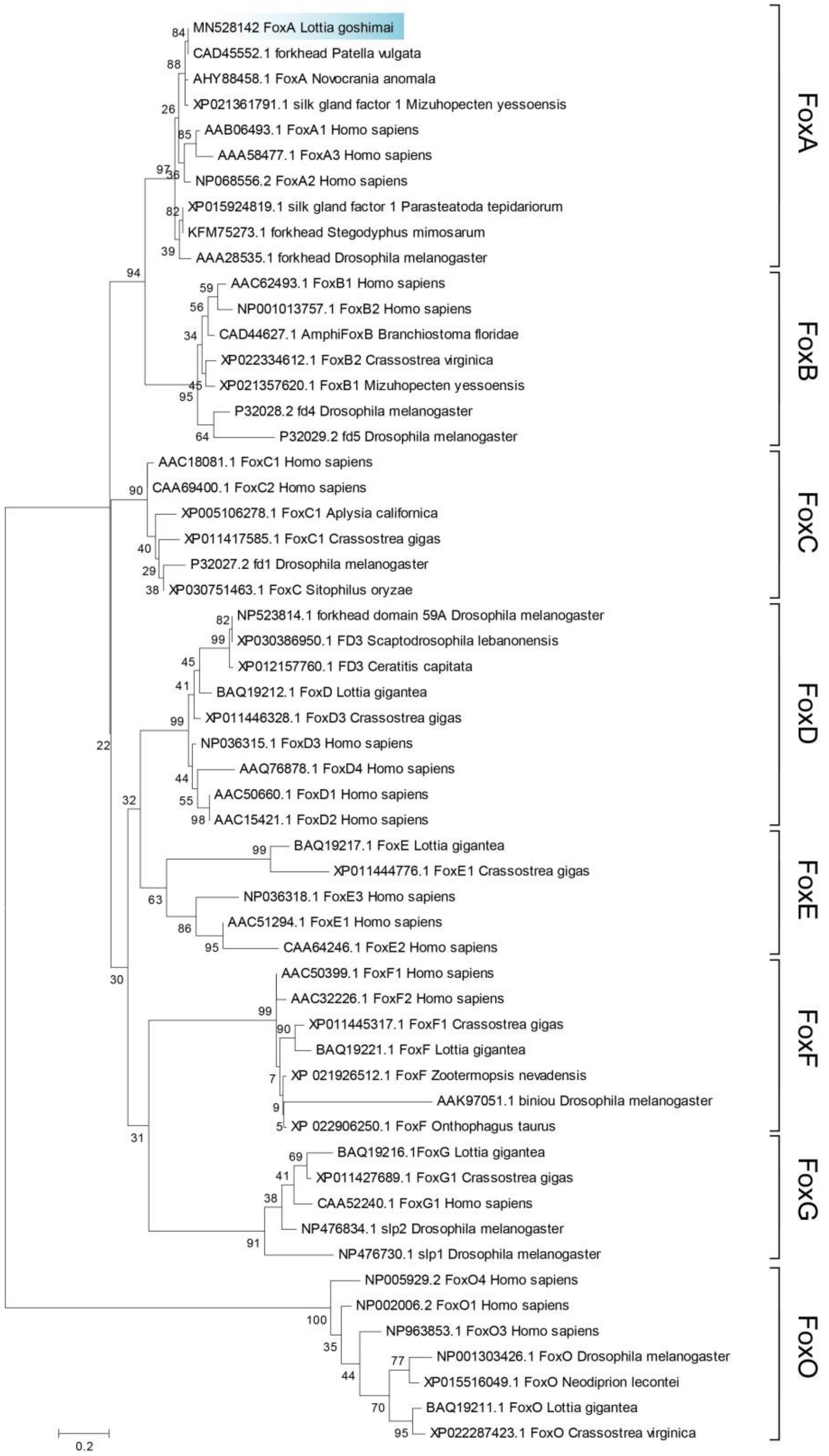


**Fig. S14 ML tree of FOX genes.** Phylogenetic analysis was performed using MEGA 6.0 with the conserved FN domain. The L + G model was estimated to be the best-fitting evolutionary model and was thus used in the analysis. The numbers at the nodes represent the bootstrap percentages from 1000 replicates.

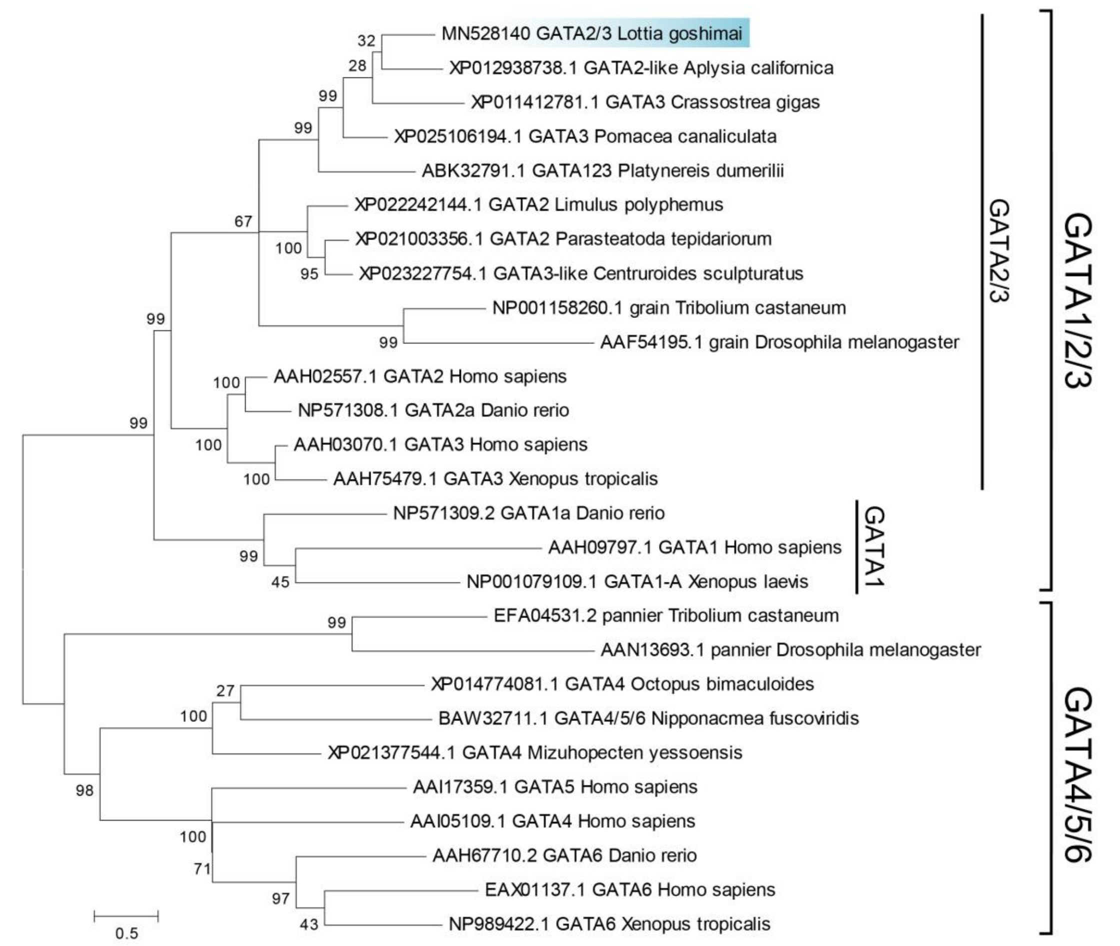


**Fig. S15 ML tree of GATA genes.** Phylogenetic analysis was performed using MEGA 6.0 with the whole amino acid sequences. The Dayhoff + G + I + F model was estimated to be the best-fitting evolutionary model and was thus used in the analysis. The numbers at the nodes represent the bootstrap percentages from 1000 replicates.

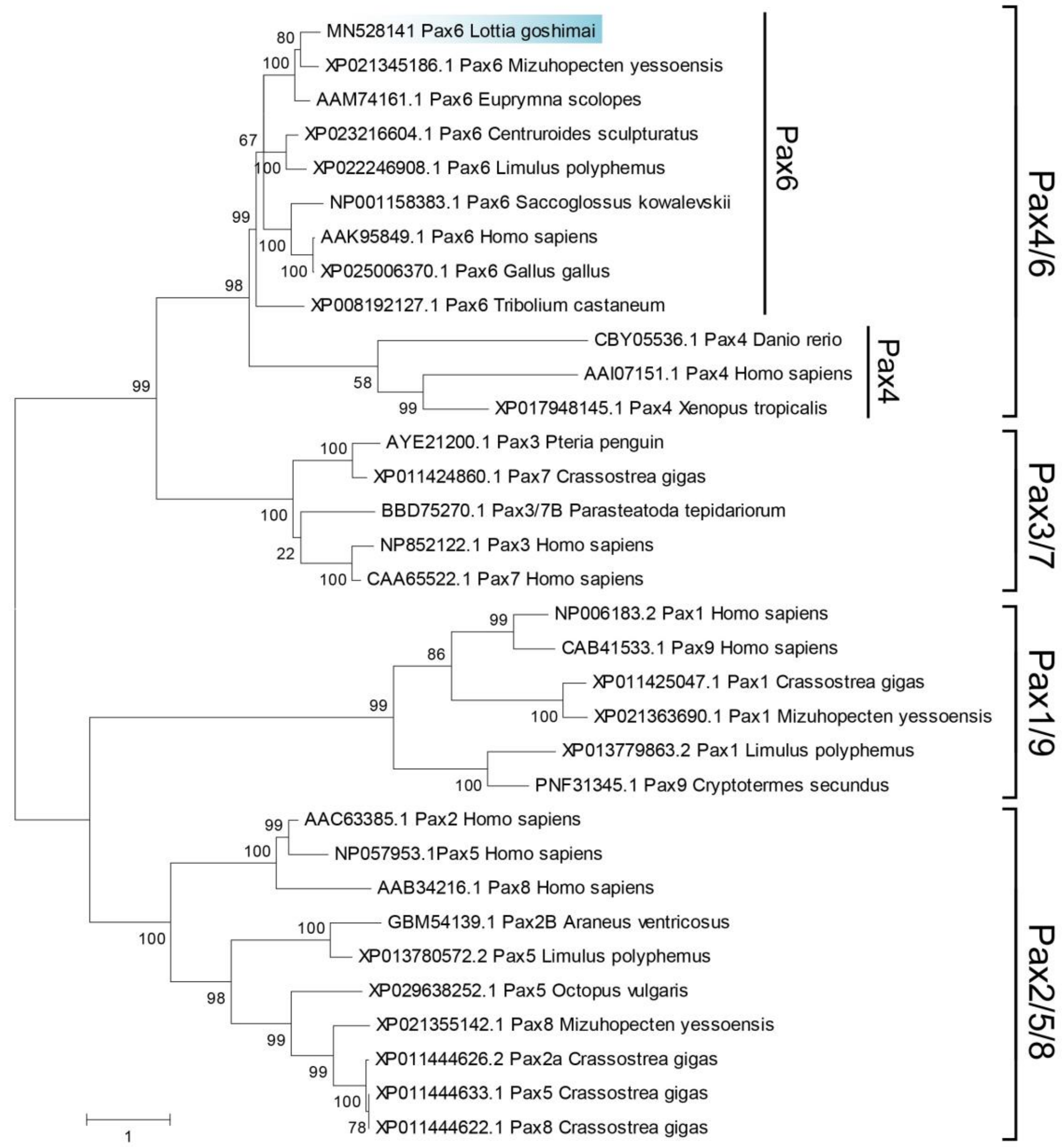


**Fig. S16 ML tree of Pax genes.** Phylogenetic analysis was performed using MEGA 6.0 with the whole amino acid sequences. The JTT + G + I + F model was estimated to be the best-fitting evolutionary model and was thus used in the analysis. The numbers at the nodes represent the bootstrap percentages from 1000 replicates.

### Caption for supplemental dataset 1

**Supplemental dataset 1.** Sequences of Chordin genes used for the phylogenetic analysis shown in supplemental Fig. S11. Some sequences were derived from the transcriptomic data kindly provided by Dr. Ferdinand Marlétaz (3).
